# A simple model for learning in volatile environments

**DOI:** 10.1101/701466

**Authors:** Payam Piray, Nathaniel D. Daw

**Affiliations:** Princeton Neuroscience Institute, Princeton University

## Abstract

Sound principles of statistical inference dictate that uncertainty shapes learning. In this work, we revisit the question of learning in volatile environments, in which both the first and second-order statistics of observations dynamically evolve over time. We propose a new model, the volatile Kalman filter (VKF), which is based on a tractable state-space model of uncertainty and extends the Kalman filter algorithm to volatile environments. The proposed model is algorithmically simple and encompasses the Kalman filter as a special case. Specifically, in addition to the error-correcting rule of Kalman filter for learning observations, the VKF learns volatility according to a second error-correcting rule. These dual updates echo and contextualize classical psychological models of learning, in particular hybrid accounts of Pearce-Hall and Rescorla-Wagner. At the computational level, compared with existing models, the VKF gives up some flexibility in the generative model to enable a more faithful approximation to exact inference. When fit to empirical data, the VKF is better behaved than alternatives and better captures human choice data in two independent datasets of probabilistic learning tasks. The proposed model provides a coherent account of learning in stable or volatile environments and has implications for decision neuroscience research.

## Introduction

Our decisions are guided by our ability to associate environmental cues with the outcomes of our chosen actions. Accordingly, a central theoretical and empirical question in behavioral psychology and neuroscience has long been how humans and other animals learn associations between cues and outcomes. According to both psychological and normative models of learning [1–4], when animals observe pairings between cues and outcomes, they update their belief about the value of the cues in proportion to prediction errors, the difference between the expected and observed outcomes. Importantly, the degree of this updating depends on a stepsize or learning rate parameter. Although some accounts take this as a free parameter, analyses based on statistical inference, such as the Kalman filter [5], instead demonstrate that the learning rate should in principle depend on the learner’s uncertainty. The dynamics of uncertainty – and hence, of learning rates – then depend on the assumed or learned dynamics of the environment. For instance, the Kalman filter is derived assuming that the true associations fluctuate randomly, but at a known, constant speed. In this case the asymptotic uncertainty, and learning rate, are determined by how quickly the associations fluctuate and how noisily they are observed. However, in volatile environments, in which the speed by which true associations change might itself be changing, uncertainty (and learning rates) should fluctuate up and down according to how quickly the environment is changing [6,7]. This normative analysis parallels classical psychological theories, such as the Pearce-Hall model [3], which posit that surprising outcomes increase the learning rate while expected ones decrease it. Those models measure surprise by the absolute value of the discrepancy between the actual outcome and the expected value, i.e. the unsigned prediction error [3].

Behavioral studies have also shown that human learning in volatile environments is consistent with the predictions of this class of models [6,8]: learning rates fluctuate with the volatility of the environment. There is also evidence for a neural substrate for such dynamic learning rates: for instance, neuroimaging studies have shown that activity in the dorsal anterior cingulate cortex covaries with the optimal learning rate [6] and recent work suggests a mechanistic model of adaptive learning rate [9]. Theoretical and empirical work also suggests that neuromodulatory systems, particularly acetylcholine, norepinephrine and serotonin, might be involved in encoding uncertainty signals necessary for computing learning rate in stable and volatile environments, respectively [7,8,10–12]. Finally, various approximate learning algorithms have been fruitful for studying individual differences in learning [13–16].

The theoretical studies establish the question of *why* the learning rate should be dynamically adjusted, and the empirical studies provide evidence that it does so. However, a complete understanding of a learning system requires understanding of *how* these theories could be realized at the process or algorithmic level [17]. This is as yet much less clear: as discussed below, statistically grounded theories of learning under volatility tend to be somewhat impractical and opaque. Furthermore, while their general analogy with the more rough and ready psychological theories like Pearce-Hall seems clear, there is not a direct mapping comparable to the way for example the Kalman filter encompasses and rationalizes the classical Rescorla-Wagner theory.

A fully Bayesian account of learning turns on two different aspects. The first is a generative model, that is a set of explicit probabilistic assumptions about how the environment evolves and generates outcomes. Inverting the generative model with Bayes’ rule gives rise to an optimal inference algorithm for estimating the latent environmental variables from observable outcomes. In situations more complicated than the smooth world of the Kalman filter, however, exact inference is generally intractable and the second aspect comes into play: additional approximations are required to achieve a practical, algorithmic- or process-level inference model. An influential model, proposed by Behrens and colleagues [6] addressed the first but not the second of these points. It comprises a two-level generative model for learning, in which a variable governing the observed associations fluctuates according to a Gaussian random walk with a speed (i.e., diffusion variance) determined, at each timepoint by a second higher-level, variable, itself fluctuating with analogous dynamics. Although this model has been conceptually influential, it lacked any tractable or biologically-plausible inference model at the algorithmic level: instead, its predictions were simulated by brute-force integration. Another model, called the hierarchical Gaussian filter (HGF) [8,18] extended Behrens’ generative model to a more general one consisting of multiple levels of hierarchy, in which the extent of diffusion noise at each level is determined by the preceding level [19]. Importantly, it also addressed the second issue by offering a tractable approximate inference rule for the process, based on a variational approach. This filter provides a biologically-plausible model for hierarchical learning and is able to capture dynamics over an arbitrary number of cascading layers with an algorithm that is easily generalizable.

Here we revisit the question of approximate inference in a two-level model of learning in volatile environments. A key theoretical complication faced by the aforementioned models is that the variables at each level of the hierarchy, above the first, represent variances for the corresponding random walks at the next level down. Since variances must be positive, if the hierarchy is taken as a cascade of analogous unbounded Gaussian random walks, nonlinear transformations must be introduced at each stage to ensure positivity. This, in turn, complicates inference: in particular, even after employing a variational approximation (as the HGF does) to decouple inference about each variable from the others, solving the resulting subproblems still requires further approximation to accommodate the nonlinearity at each stage. Informed by this reasoning, we propose a novel model for learning in volatile environments that is conceptually derived from that of Behrens et al. [6] and the 2-level case of the HGF. However, we introduce a distinct diffusion dynamics for the upper level, which ensures positivity and also an exact, conjugate solution to the variational maximization. The resulting model thus gives up some of the elegant flexibility of the HGF (its ability recursively to chain through an arbitrarily deep hierarchy) in return for a simpler inference rule requiring fewer approximations for the most widely used, 2-level, case. We separately encounter, and take a distinct approach to, a second issue of nonlinearity that arises in these models, which arises when (as has almost always been the case in empirical studies of human learning in this area) the observable outcomes like rewards are binary-valued instead of continuous. The resulting algorithm, called VKF, is a generalization of Kalman filter algorithm to volatile environments and resembles models that hybridize the error-driven learning from the Rescorla-Wagner model and the Kalman filter with Pearce-Hall’s dynamic learning rate (as proposed by different authors, for example by Li et al. and Le Pelley [20,21]). Notably, in volatile environments, the learning rate fluctuates with larger and smaller than expected prediction errors, as suggested by models such as Pearce-Hall.

In the next section, we review the Kalman filter algorithm and present the generative model underlying the VKF and the resulting learning algorithm. The full formal treatment is given in the S1 Appendix. Next, we show that the proposed model outperforms existing models in predicting empirical data.

## Results

### Theoretical results

#### Kalman filter for tracking in environments with stable dynamics

The Kalman filter is the cornerstone of statistical tracking theories, with widespread applications in many technological and scientific domains including psychology and neuroscience [4,7,22,23]. The Kalman filter corresponds to optimal statistical inference for a particular class of linear state space environments with Gaussian dynamics. In those environments, the hidden state of the environment is gradually changing across time (according to Gaussian diffusion) and the learner receives an outcome on each time depending on the current value of the state (plus Gaussian noise). In these circumstances, the posterior distribution over the hidden state is itself Gaussian, thus tracking it amounts to maintaining two summary statistics: a mean and a variance.

Formally, consider the simplest case of prediction: that of tracking a noisy, fluctuating reward (e.g. that associated with a particular cue or action), whose magnitude *o*_*t*_ is observed on each trial *t*. Assume a state space model in which, on trial *t*, the hidden state of the environment, *x*_*t*_ (the true mean reward), is equal to its previous state *x*_*t*−1_ plus some process noise

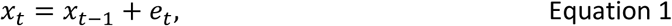

where the process noise *e*_*t*_∼*Normal*(0, *v*) has a variance given by *v*. A critical assumption of the Kalman filter is that the process uncertainty, *v*, is constant and known. The outcome then noisily reflects the hidden state, *o*_*t*_∼*Normal*(*x*_*t*_, *σ*^2^). Here the observation noise is again Gaussian with known, fixed variance, *σ*^2^. The Kalman filtering theory indicates that the posterior state at time *t*, given all previous observations, *o*_1_, …, *o*_*t*_, will itself be Gaussian. Its mean, *m*_*t*_, is updated at each step according to the prediction error:

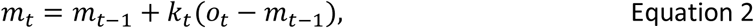

where *k*_*t*_ is the learning rate (or Kalman gain)

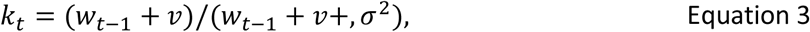

which depends on the noise parameters and the posterior variance *w*_*t*−1_ on the previous trial. Note that *k*_*t*_ < 1 in all trials and it is larger for larger values of *v*. On every trial, the posterior variance also gets updated:

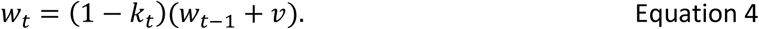

Note that although it is not required for prediction (that is for updating *m*_*t*_), it is also possible to compute the autocovariance (given observations *o*_1_,…, *o*_*t*_), defined as the covariance between consecutive states, *w*_*t*−1,*t*_ = cov[*x*_*t*−1_, *x*_*t*_], which is given by

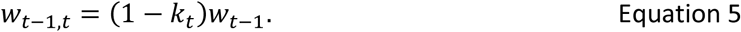

This equation indicates that when the Kalman gain is relatively small, the autocovariance is large, which means that information transmitted by observing a new outcome is expected to be quite small. We will see in the next section that this autocovariance plays an important role in inference in volatile environments.

#### VKF: A novel algorithm for tracking in volatile environments

We next consider a volatile environment in which the dynamics of the environment might themselves change. In the language of the state space model presented above, the process noise dynamically changes. Thus, the variance of the process noise (*e*_*t*_ in Equation 1) is a stochastic variable changing with time (Figure 1A). To build a generative model, we need to make some assumptions about the dynamics of this variable. Our approach here is essentially the same as that taken by Smith and Miller [24] and by Gamerman et al. [25] (see also West et al. [26]). Consider a problem in which the process variance dynamically changes. In the previous section, we saw that the state *x*_*t*_ diffused according to additive noise. Because variances are constrained to be positive, it makes sense to instead assume their diffusion noise is multiplicative to preserve this invariant. Therefore, we assume that the current value of precision (inverse variance), *z*_*t*_, is given by its previous value multiplied by some independent noise.

**Fig 1.**
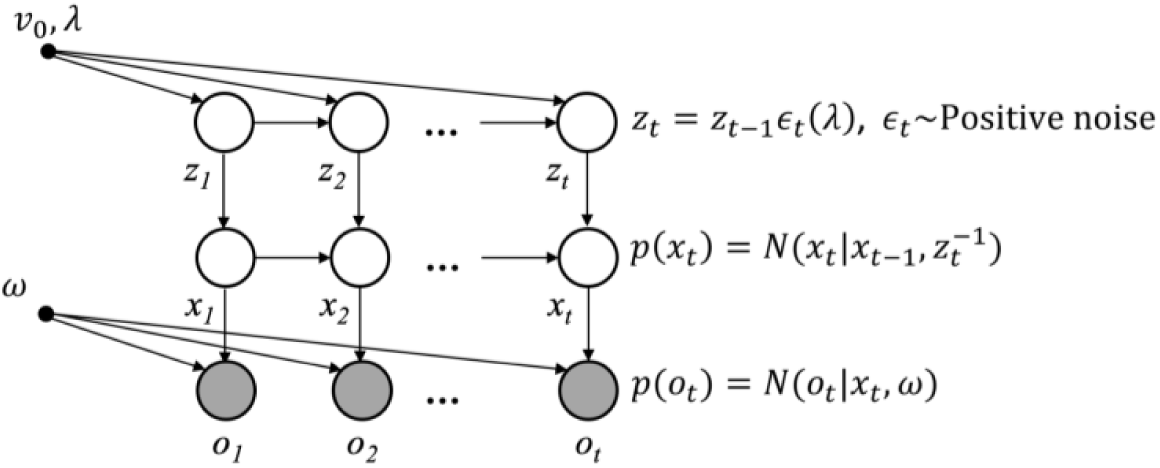
The generative model of VKF. The generative model consists of two interconnected hidden temporal chains, *x*_*t*_ and *z*_*t*_, governing observed outcomes, *o*_*t*_. Arrows indicate the direction of influence. On each trial, *t*, outcome, *o*_*t*_, is generated based on a Gaussian distribution, its mean is given by the hidden random variable *x*_*t*_, and its variance is given by the constant parameter *ω*. This variable is itself generated according to another Gaussian distribution, which its mean is given by *x*_*t*−1_, and its variance is given by another hidden random variable, 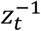. This variable is itself generated based on its value on the previous trial, *z*_*t*−1_, multiplied by some positive noise distributed according to a scaled Beta distribution governed by the parameter *λ*. The inverse mean of this variable on the first trial is assumed to be given by another constant parameter, *v*_0_. See Equations 1 and 6 for further explanation.

Formally, the current state of *z*_*t*_ is given by

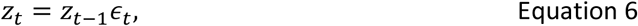

where *ϵ*_*t*_ is an independent random variable on trial *t*, which is distributed according to a rescaled beta distribution (as detailed in S1 Appendix), such that the mean of *ϵ*_*t*_ is 1 (that is, conditional expectation of *z*_*t*_ is equal to *z*_*t*−1_) but has spread controlled by a free parameter 0 < *λ* < 1. The value of noise, *ϵ*_*t*_, is always positive and is smaller than (1 − *λ*)^−1^. Therefore, while “on average”, *z*_*t*_ is equal to *z*_*t*−1_, *z*_*t*_ can be smaller than *z*_*t*−1_, or larger by a factor up to (1 − *λ*)^−1^. Thus, the higher the parameter *λ*, the faster the diffusion.

To build up the full model, first consider a simplified case in which the latent state 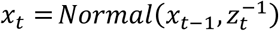 is directly observed at each step and, further, has some known mean *μ*_*t*_. Although this sub-problem greatly simplifies the main problem by isolating *x*_*t*_ from *x*_*t*−1_, it provides the foundation for inference in the full model. This is because it is similar to the problem that remains once a variational approximation is introduced. It can be formally shown (see S1 Appendix) that in the simplified case, estimating the process variance, i.e. the posterior over *z*_*t*_ at each step given all previous observations, is tractable. Specifically, the posterior distribution over *z*_*t*_ takes the form of a Gamma distribution, whose inverse mean, *v*_*t*_, is updated according to the observed sample variance:

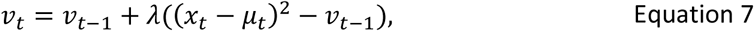

where *v*_*t*_ = *E*[*z*_*t*_]^−1^. Note that this equation amounts to an error-correcting rule for updating *v*_*t*_, in which the error is given by (*x*_*t*_ − *μ*_*t*_)^2^ − *v*_*t*−1_: the difference between the observed and expected squared prediction errors.

Now we are ready to build the full generative model in which both *z*_*t*_ and *x*_*t*_ dynamically evolve over time (Fig 1). In this model, as before, the observation, *o*_*t*_∼*Normal*(*x*_*t*_,*σ*^2^), follows a Gaussian distribution with the mean given by the hidden variable *x*_*t*_, which itself is given by 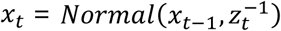: a diffusion whose precision (i.e. inverse variance) is given by another dynamic random variable, *z*_*t*_. The value of *z*_*t*_ in turn depends on its previous value multiplied by some noise that finally depends on parameter *λ* according to Equation 6. Therefore, this generative model consists of two chains of random variables, which are hierarchically connected to each other. Unlike each of those two chains separately, however, when they are conjoined in this hierarchical model, inference is not tractable and therefore we need some approximation. We use structured variational inference for approximate inference in this model [27,28]. This technique assumes a factorized approximate posterior distribution and minimizes the mismatch between this approximate posterior and the true posterior using the principle of variational inference.

The resulting learning algorithm is very similar to the Kalman filtering algorithm and encompasses Kalman filter as a special case. Importantly, the new algorithm also tracks volatility on every trial, denoted by *v*_*t*_, which is defined as the inverse of expected value of *z*_*t*_. Therefore, we call the new algorithm the “volatile Kalman filter”. In this algorithm, the update rule for the posterior mean *m*_*t*_ and variance *w*_*t*_ over *x*_*t*_ (Equations 9-13 below) is exactly the same as the Kalman filtering algorithm, but in which the constant process variance is replaced by the estimated volatility on the previous trial, *v*_*t*−1_. The volatility also gets updated on every trial according to expected value of (*x*_*t*_ − *x*_*t*−1_)^2^

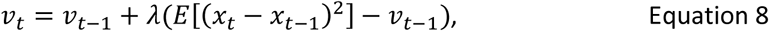

where the expectation should be taken under the approximate posterior over *x*_*t*−1_ and *x*_*t*_. Therefore, the volatility update rule takes a form of error correcting, in which the error is given by *E*[(*x*_*t*_ − *x*_*t*−1_)^2^] − *v*_*t*−1_ with the noise parameter, *λ*, as the step size. Thus, the higher the noise parameter, the higher the speed of volatility update is. Therefore, we call *λ* the “volatility update rate”. Also, note that the expectation in this equation depends on the autocovariance.

It is then possible to write *E*[(*x*_*t*_ − *x*_*t*−1_)^2^] in terms of the variance and covariance of *x*_*t*−1_ and *x*_*t*_ to obtain Equation 8 below and complete the VKF learning algorithm:

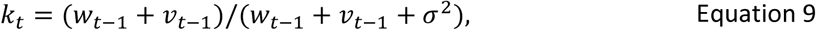

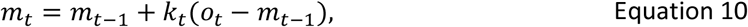

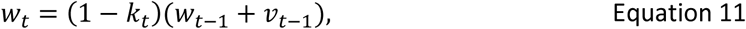

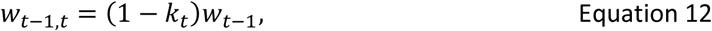

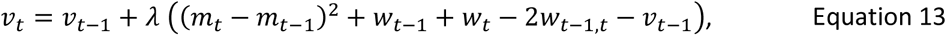

where *σ* is the constant variance of observation noise. In addition to the volatility update parameter, *λ*, which is constrained in the unit range, this algorithm depends on another parameter, *v*_0_ > 0, which is the initial value of volatility. Notably, the Kalman filter algorithm is a special case of the VKF in which *λ* = 0 and the process variance is equal to *v*_0_ on all trials. In the next section, we test this model with synthetic and empirical datasets.

#### Binary VKF

We have also developed a binary version of VKF for situations in which observations are binary. The generative model of the binary VKF is the same as that of VKF with the only difference that binary outcomes are generated according to Bernoulli distribution with the parameter given by *s*(*x*_*t*_) = 1/(1 + exp(−*x*_*t*_)), where *s*(*x*_*t*_) is the sigmoid function, which maps the normally distributed variable *x*_*t*_ to the unit range. For this generative model, the inference is more difficult because the relationship between hidden states and observations is nonlinear. Therefore, further approximation is required to perform inference here, because observations are not normally distributed and Equation 1 does not hold. For the binary VKF, we assumed a constant posterior variance, *ω*, and employed moment matching (which is sometimes called assumed density filtering [29,30]) to obtain the posterior mean. The resulting algorithm is then very similar to the original VKF with the only difference that the update rule for the mean (i.e. Equation 10) is slightly different:

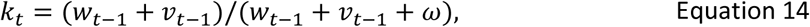

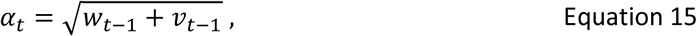

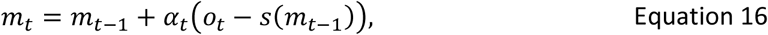

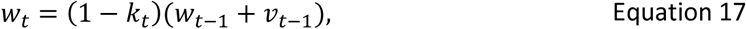

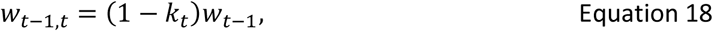

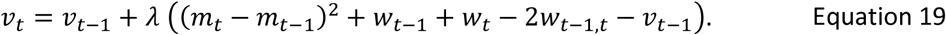

Note that the learning rate for the binary VKF is *α*_*t*_ defined by Equation 15. Furthermore, we have introduced a parameter, *ω* > 0, specifically for inference (i.e. does not exist in the generative model). We call this parameter the noise parameter, because its effects on volatility are similar to the noise parameter, *σ* for linear observations (through *k*_*t*_, Equation 14). However, it is important to note that unlike *σ*, this parameter does not have an inverse relationship with the learning rate, *α*_*t*_.

### Simulation analyses

#### Comparing VKF with known ground truth

First, we study the performance of VKF on simulated data with known ground truth. We applied the VKF to a typical volatility tracking task (similar to [6,8]). In this task, observations are drawn from a normal distribution whose mean is given by the hidden state of the environment. The hidden state is constant +1 or –1. Critically, the hidden state was reversed occasionally. The frequency of such reversals itself changes over time; therefore, the task consists of blocks of stable and volatile conditions. We investigate performance of VKF in this task because its variants have been used for studying volatility learning in humans [6,8]. Note that, as here, it is common to study these models’ learning performance when applied to situations that do not exactly correspond to the generative statistical model for which they were designed. In particular, it has been shown using this type of task that humans’ learning rates are higher in the volatile condition [6,8]. Fig 2 shows the performance of VKF in this task. As this figure shows, the VKF tracks the hidden state very well and its learning rate is higher in the volatile condition. Furthermore, the volatility signal increases when there is a dramatic change in the environment. Note that as for other models previously fit to tasks of this sort, the switching dynamics at both levels of hidden state here are not the same as the random walk generative dynamics assumed by our model. These substitutions demonstrate that the principles relating volatility to learning are general, and so the resulting models are robust to this sort of variation in the true generative dynamics.

**Fig 2.**
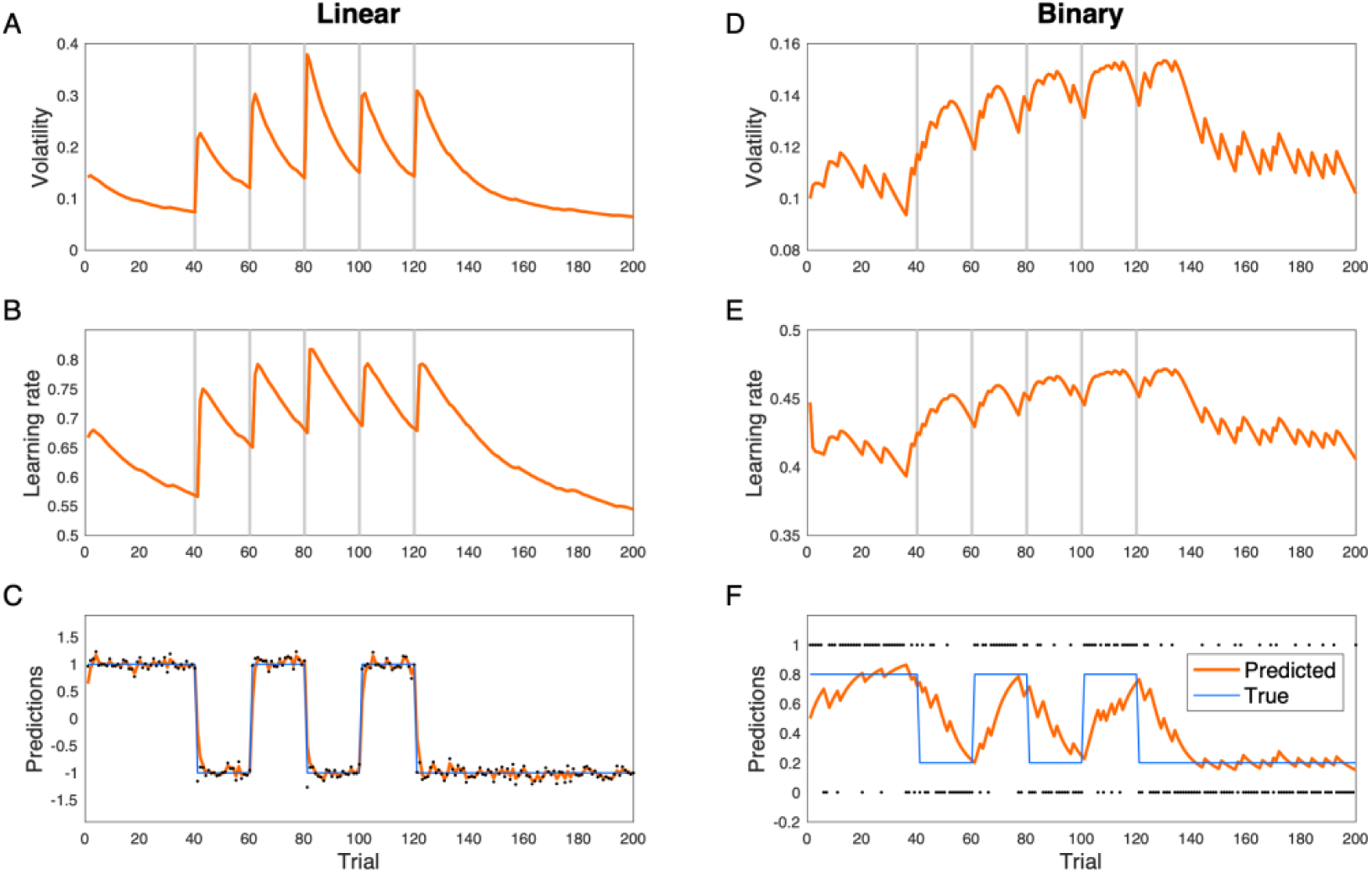
Behavior of the VKF. A-C) A switching probabilistic learning task in which observations are randomly drawn from a hidden state (with variance 0.01) that switches several times. The relationship between the hidden state and observations are either linear (A-C) or binary (D-F). The volatility signal with the VKF increases after a switch in the underlying hidden state and the learning rate signal closely follows the volatility. The grey lines show the switching time. Dots are actual observations. These parameters used for simulating VKF: *λ* = 0.1, *v*_0_ = 0.1, *σ* = 0.1, *ω* = 0.1.

In another simulation analysis, we studied the performance of the binary VKF in a similar task, but now with binary observations, which were used in previous studies [6,8]. Here, observations are drawn from a Bernoulli distribution whose mean is given by the hidden state of the environment. The hidden state is a constant probability 0.8 or 0.2, except that it is reversed occasionally. As Fig 2 shows, predictions of the binary VKF match with the hidden state. Furthermore, the volatility signal increases when there is a dramatic change in the environment, and the learning rate is higher in the volatile condition. Similar simulation analyses with higher levels of volatility show the same behavior: the volatility signal and the learning rate are generally larger in volatile blocks, in which the environment frequently switches (Fig 3).

**Fig 3.**
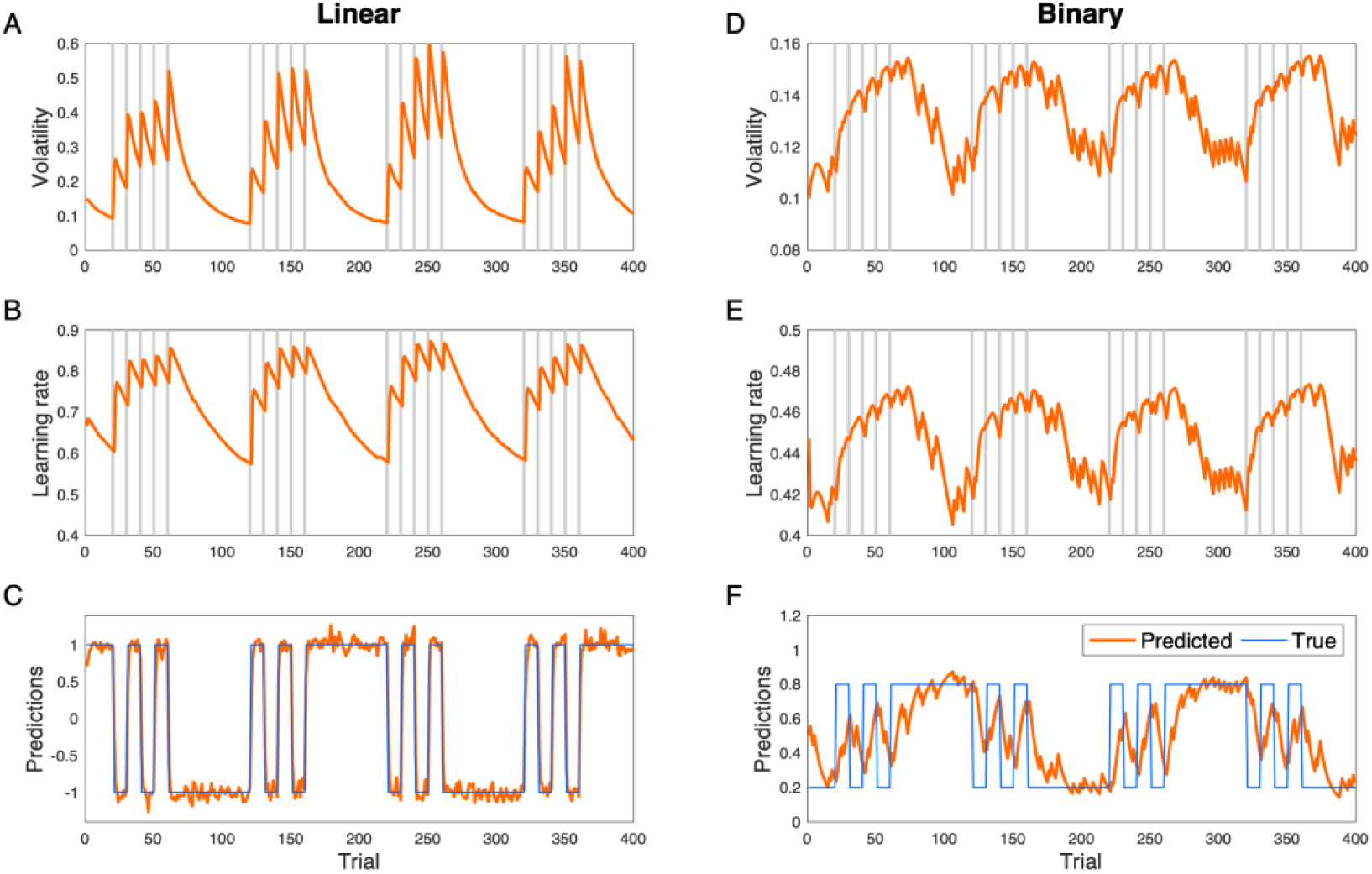
Behavior of the VKF in highly volatile condition for linear. (A-C) and binary (D-F) observations. The volatility and learning rate signal are larger in volatile conditions in which the hidden state is frequently changing. The parameters used were the same as Fig 2.

The VKF is a relatively simple model that approximates exact solution to an intractable inference problem. To further address the question how accurate is this approximation, next, we compared the performance of the VKF to other approaches: first, one representing (as close as was feasible) exact Bayesian inference, and, second, a different approach to approximate inference, the HGF.

#### Comparing VKF with the particle filter

We first compared the behavior of the VKF with a computationally expensive but near-exact method as the benchmark. For sequential data, the particle filter is a well-known Monte Carlo sequential sampling method, which approximates the posterior at every timepoint with an ensemble of samples; it approaches optimality in the limit of infinite samples. We used a Rao-Blackwellized particle filter (RBPF) [31] for this analysis, which combines sampling of some variables with analytical marginalization of others, conditional on these samples. In this way, it exploits the fact that inference on the lower level variable (i.e. *x*_*t*_ in Fig 1) is tractable using a Kalman filter, given samples from the upper level variable (i.e. *z*_*t*_ in Fig 1). Therefore, this approach essentially combines particle filter with Kalman filter. We used the same sequence generated in previous analyses (Fig 2) and compared the RBPF algorithm with 10,000 samples as the benchmark, assuming the same parameters for both algorithms. As shown in Fig 4, the behavior of VKF is very well matched with that of this benchmark for Gaussian observations. In particular, predictions and volatility estimates of the two algorithms were highly correlated, with correlation coefficients of 1.00 and 0.95, respectively, in this task. We also quantified the relative error, defined as the average mismatch between predictions of VKF and ground truth lower-level states, *x*_*t*_, measured in units of increased error relative to the benchmark error of RBPF (see Methods), as 22.9%.

**Fig 4.**
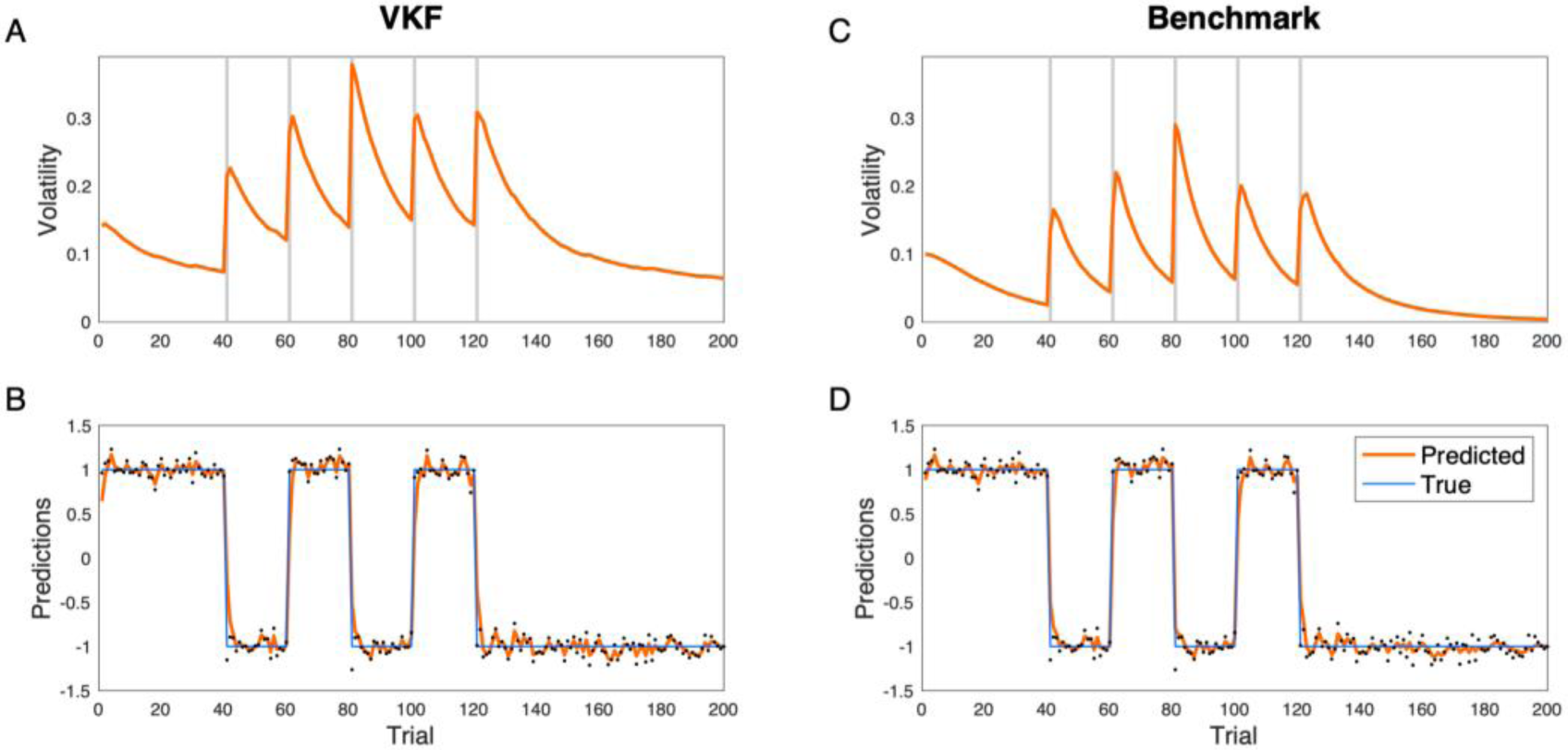
Comparison of VKF with the benchmark sampling methods. RBPF was used as the benchmark (with 10000 particles). The behavior of the VKF closely follows that of the benchmark. The parameters used were the same as Fig 2.

We next compared the performance of the binary VKF with that of the particle filter benchmark. For the binary VKF, the inference is more difficult because the relation between the state and the observation is also nonlinear. For the same reason, it is not possible to use RBPF to marginalize part of the computation here because the submodel conditional on the variance level is also not analytically tractable; instead we used a conventional particle filter sampling across both levels. We ran the particle filter on this problem with 10,000 samples and the same parameters as that of the binary VKF. As Fig 5 shows, the latent variables estimated by the VKF and the benchmark particle filter are again quite well matched, although the particle filter responds more sharply following contingency switches. Similar to the previous analysis, predictions and volatility estimates were well correlated with correlation coefficients given by 0.91 and 0.68, respectively. The relative error in state estimation for binary VKF was 5.4%.

**Fig 5.**
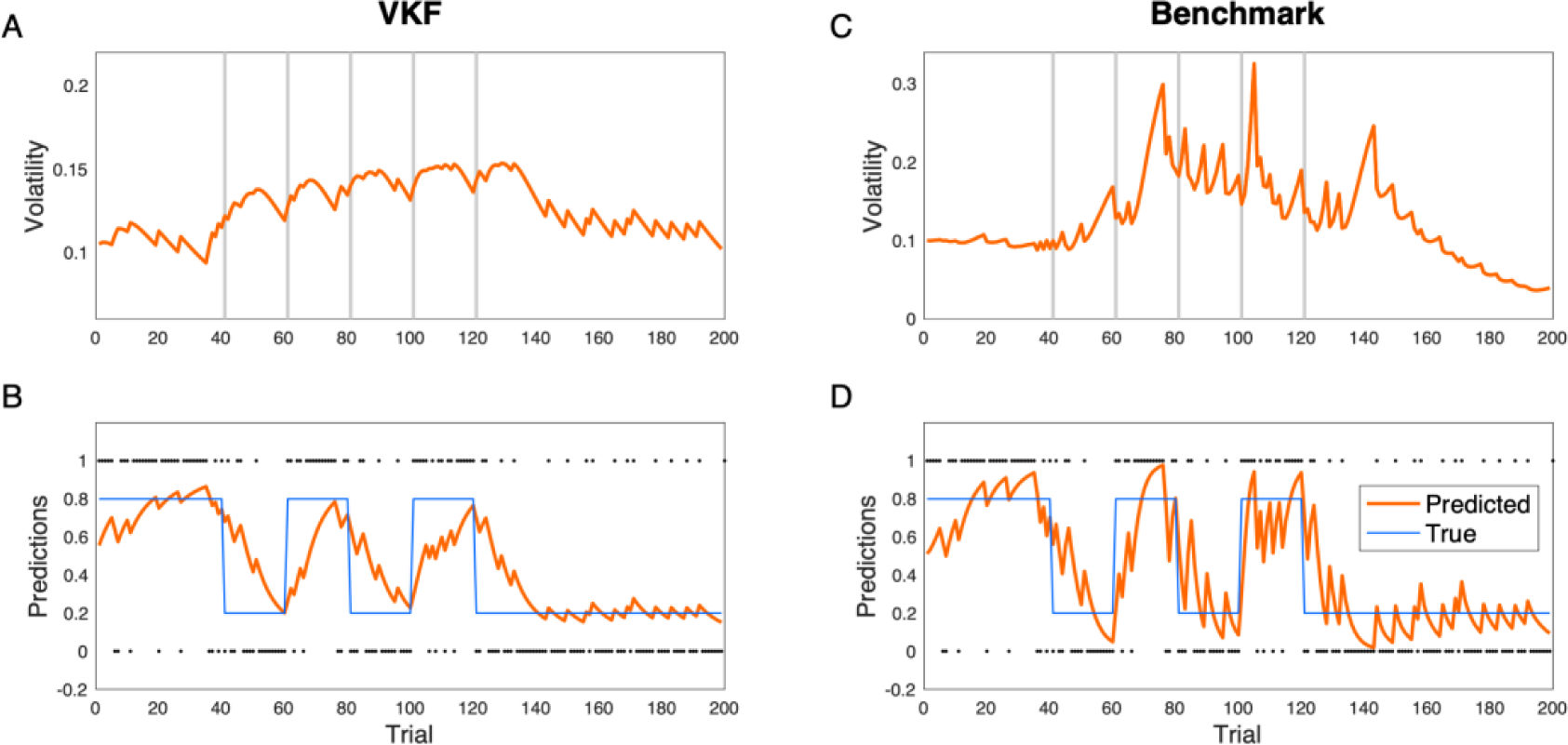
Comparison of binary VKF with the benchmark sampling methods. Particle filter was used as the benchmark (with 10000 particles). Both the VKF and the benchmark show higher learning rate in the volatile condition and estimated volatility signals by the two models are highly correlated. The parameters used were the same as Fig 2.

#### Comparing VKF with HGF

We next compared VKF to another approximate approach with more comparable computational complexity. In particular, we compared it to the HGF (with two latent levels), as the latter is the most commonly used algorithm for learning in volatile environments. Similar to our model, the generative model of the HGF assumes that there are two chains of state variables, *x*_*t*_ and *z*_*t*_, in which the distribution over *x*_*t*_ is given by a normal distribution with the mean and variance depending on *x*_*t*−1_ and *z*_*t*_, respectively. These two algorithms differ in two ways, however. First, for the generative models, the form of the process noise controlling the diffusion of *z*_*t*_ is different. In our model, the noise is a positive variable with a beta distribution. The HGF, however, assumes that the process noise for *z*_*t*_ is additive and normally-distributed, and hence exponential transformation is needed to ensure that the variance of *x*_*t*_ is non-negative. These differences, in turn, give rise to differences in the resulting approximate inference rules, particularly for inference over the volatility level, *z*_*t*_. Our model has specific conjugacy properties, which make the inference for *z*_*t*_ tractable if *x*_*t*_ is isolated from *x*_*t*−1_ (e.g. if *x*_*t*−1_ is fixed and under a variational approximation; see theoretical explanation above and S1 Appendix for mathematical proofs). However, even in this case when *x*_*t*_ is isolated from *x*_*t*−1_, the inference over *z*_*t*_ in the HGF is not tractable and therefore requires further approximation. The approximate inference model of the HGF thus relies on a second-order Taylor approximation to deal with this issue, in addition to the variational approximation used by both the VKF and the HGF. By eliminating this additional approximation, the VKF avoids one source of potential inaccuracy. Also, the VKF’s conjugacy ensures a simple (one-parameter Gamma) distribution for the posterior estimate at the top (volatility) level, which may also contribute to stability, vs. a two-parameter Gaussian approximation, in which numerical problems (e.g., negative values for the approximated posterior uncertainty) can arise.

However, directly comparing the effect of these two approaches to inference is difficult because they are closely linked with two different sets of generative assumptions. To focus our comparison on the approximations, we studied the ability of each algorithm to reproduce data generated from its own, corresponding generative model, in roughly comparable parameter regimes. Then to ensure any differences were related to the approximate inference itself, rather than to the different generative statistics themselves (e.g., supposing one problem is just harder), we scored each model in terms of error relative to optimal inference as proxied by the particle filter. This represents the best achievable performance for each specific generative process. We also performed a number of followup analyses to further pursue this question, in S1 Text.

Thus, we generated a sequence of observations using the generative model of the HGF (see Methods). We then used the generated sequence as the input to the HGF inference algorithm, using true generative parameters. We repeated this process 1000 times. In 82 of these simulations, the HGF encountered numerical problems: its inferred trajectory encountered numerical problems, i.e. negative estimates of the posterior variance over the top (volatility) level. This problem is due to the Taylor approximation used to extrapolate the variational posterior in the approximate inference model of the HGF. Only for the remaining 918 simulations were we able to quantify the error of HGF, defined as the mismatch between predicted and true lower level states *x*_*t*_ (see Methods). We also performed inference on the same generated data using a RBPF (derived under the HGF generative assumptions) [31], as a proxy of exact inference. We compared these two results to obtain a measure of fractional relative error in the HGF over and above the (unavoidable) error from the RBPF. Similarly, we quantified the error of the VKF relative to the RBPF: data were simulated under the VKF generative model (see Methods), which were then used as the input to the VKF inference model and its associated RBPF. This analysis revealed that the relative error of the HGF and VKF is 21.4% (SE=6.8%) and 2.7% (SE=0.3%), respectively, indicating that the VKF performance is closer to the particle filter than the HGF. We performed a number of control analyses confirming these results with different sets of parameters, and using an alternative baseline independent of the RBPF (S1 Text).

### Effects of volatility on learning rate for binary outcomes

Following Behrens’ seminal study [6], the majority of previous work studying volatility estimation used binary outcomes. This is also important from a psychological perspective, because seminal models such as Pearce-Hall indicate that learning rate, or as it is called in that literature “associability”, reflects the extent that the cue has been surprising in the past. In fact, modern accounts of learning in decision neuroscience are partly built on this classic psychological idea, and there is evidence that volatility-induced surprising events increase the learning rate in probabilistic learning tasks in humans [6,32–34]. Here, we highlight a crucial difference between the binary versions of the HGF and VKF regarding the relationship between volatility and learning rate.

We compared performance of HGF and VKF for binary observations, mimicking the sorts of experiments typically used to study volatility learning in the lab [6]. In this case, the generative dynamics of the latent variables did not match either model, but instead used a discrete change-point dynamics again derived from these empirical studies. The binary versions of the HGF and VKF use different approximations to deal with the nonlinear mapping between hidden states and binary observations, on top of the ones discussed before. Whereas the HGF employs another Taylor approximation to deal with binary observations, the binary VKF is based on a moment-matching approximation. Importantly, these different treatments of binary observations result in qualitative differences in the relationship between the learning rate and volatility. Accordingly, in the binary VKF, the learning rate closely follows the volatility estimate, similar to the VKF for continuous-valued observations and echoing the intuition from the psychological and decision neuroscience literatures. In the binary version of HGF, however, the relationship between the learning rate and volatility is more involved because the learning rate is also affected by the sigmoid transformation of the estimated mean on the previous trial. Fig 6 illustrates these signals in an example probabilistic switching task for both models. To demonstrate this point quantitatively, we generated 500 time-series and fitted both models to these time-series. The correlation between the learning rate and volatility estimates for these time-series were then calculated using the fitted parameters of each model. As expected, the learning rate and volatility signals of the binary VKF were positively correlated in all simulations (average correlation coefficient about 1.00), whereas those of binary HGF were, unexpectedly, negatively correlated in all simulations (average correlation coefficient = –0.74). Note that this qualitative difference is a general behavior of these models and is not due to a particular setting of parameters (S1 Text).

**Fig 6.**
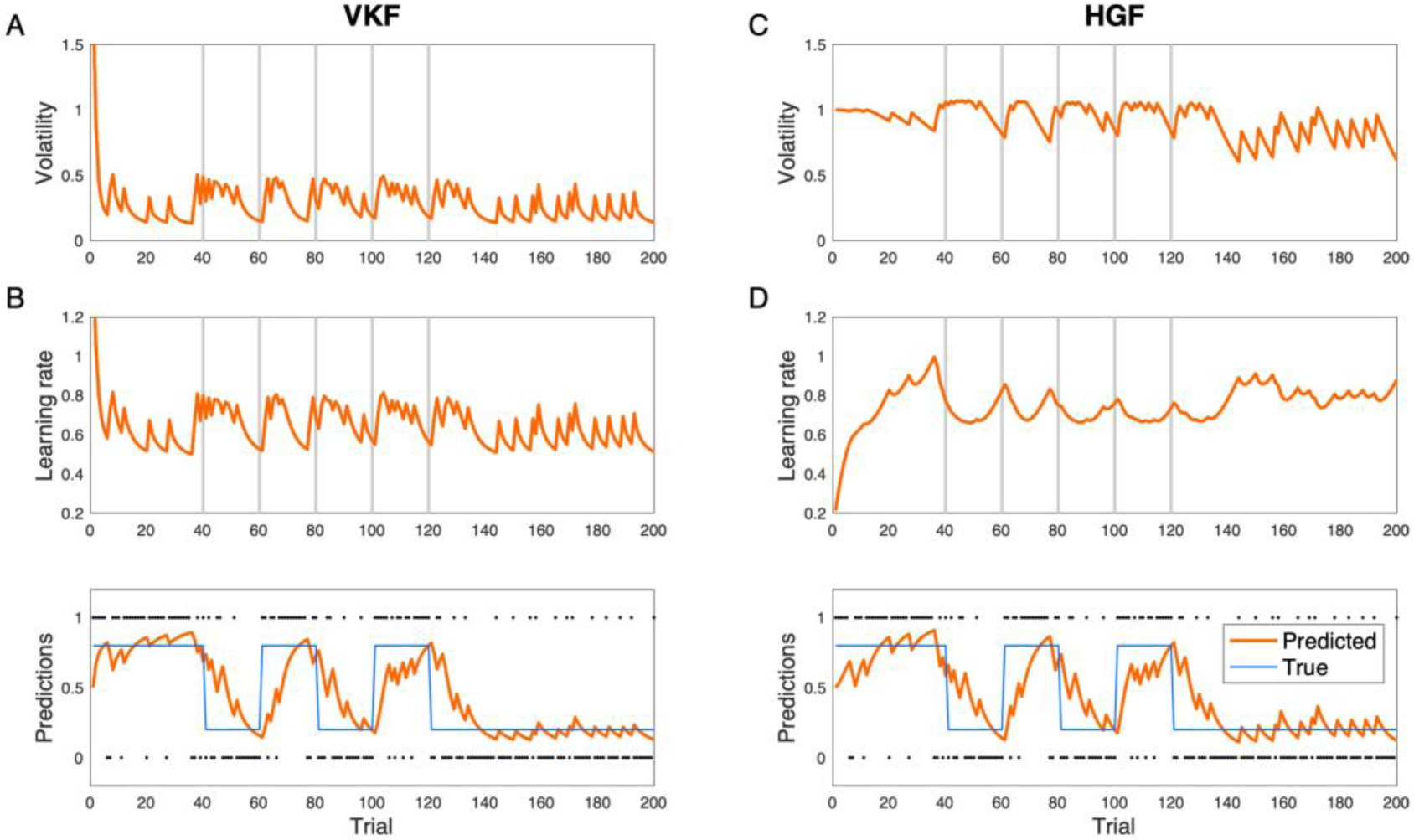
Behavior of the binary VKF and HGF. Predictions of the two models about the relationship between volatility and learning rate are different. Fitted parameters for each model were used for obtaining volatility, learning rate and prediction signals here.

### Testing VKF using empirical data

We then tested the explanatory power of the VKF to account for human data using two experimental datasets. In both experiments, human subjects performed a decision-making task, repeatedly choosing between two options, where only one of them was correct. Participants received binary feedback on every trial indicating which option was the correct one on that trial.

Such a test requires fitting the binary VKF to choice data by estimating its free parameters. The binary VKF relies on three free parameters: volatility learning rate, *λ*, the initial volatility value, *v*_0_, and the noise parameter for binary outcomes, *ω*. Fig 7 shows the behavior of the binary VKF as a function of these parameters. As Fig 7 shows, the volatility learning rate parameter determines the degree by which the volatility signal is updated and its effects are particularly salient following a contingency change. The effects of initial volatility are more prominent in earlier trials. The noise parameter indicates the scale of volatility throughout the task. Note that the noise parameter also has a net positive relationship with the learning rate in the binary VKF.

**Fig 7.**
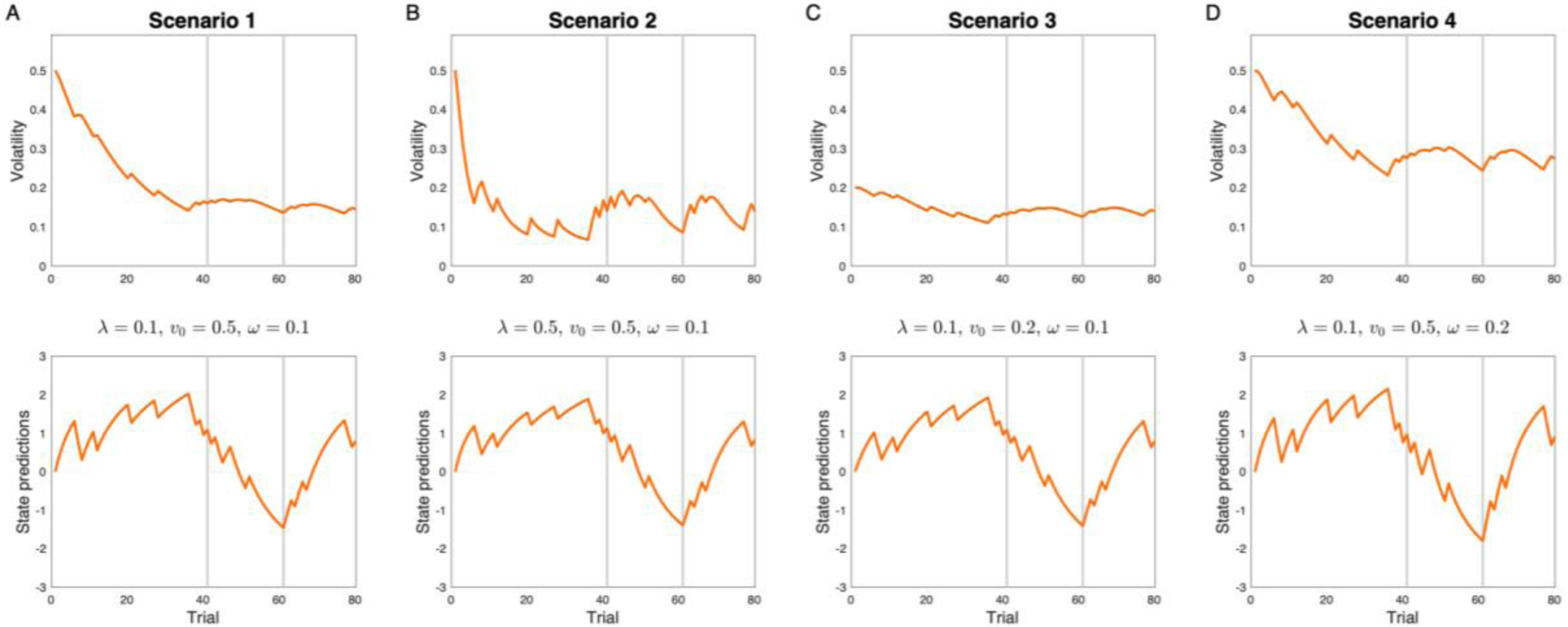
Effects of parameters on the VKF. A) The volatility and state prediction signals within the binary VKF are shown for a baseline scenario. B-D) The impact of changing parameters with respect to the baseline scenario is shown for (B) the volatility learning rate, *λ*, (C) the initial volatility *v*_0_, and (D) the noise parameter, *ω*.

To model choice data, the predictions of the binary VKF were fed to a softmax function, which has a decision noise parameter, *β*. We first verified that these parameters can be reliably recovered using simulation analyses with synthetic datasets, in which observations and choices of 50 artificial subjects were generated based on the binary VKF and the softmax (see Methods for details of this analysis). We then fitted the parameters of the VKF to each dataset using hierarchical Bayesian inference (HBI) procedure [35], an empirical Bayes approach with the advantage that fits to individual subjects are constrained according to the group-level statistics. We repeated this simulation analysis 500 times. As reported in Table 1, this analysis revealed that the model parameters were fairly well recoverable, although estimation of the volatility learning rate and initial volatility were more prone to error.

**Table 1.**
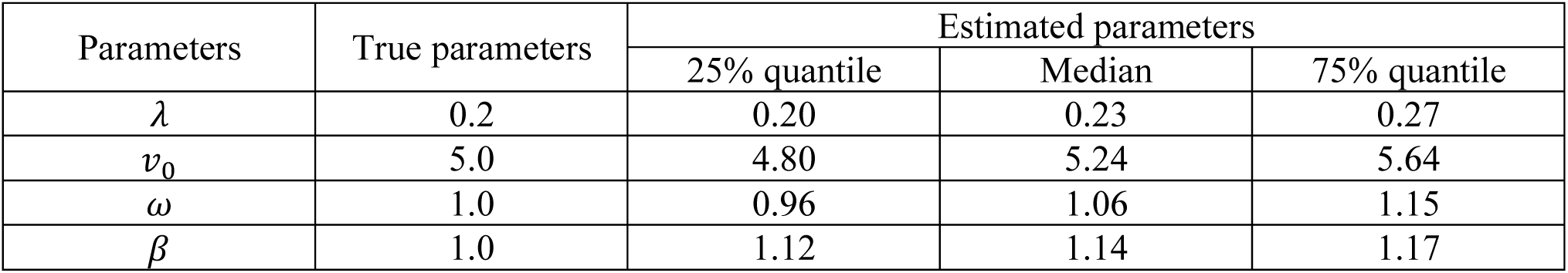
Recovery analysis for parameters of the VKF. A dataset including 50 artificial subjects were generated based on the binary VKF and a softmax choice model. The same procedure used in analysis of empirical data (HBI) was used then to estimate the parameters. We have reported the mean of parameters across all 50 artificial subjects. This procedure was repeated 500 times.

In the first experiment (Fig 8), 44 participants carried out a probabilistic learning task (originally published in [36]), in which they were presented with facial cues and were asked to make either a go- or a no-go-response (i.e., press a button, or withhold a press, respectively) for each of four facial cues in order to obtain monetary reward or avoid monetary punishment. The cues were four combinations of the emotional content of the face image (happy or angry) and its background color (grey or yellow) representing valence of the outcome. Participants were instructed that the correct response is contingent on these cues. The response-outcome contingencies for the cues were probabilistic and manipulated independently, and reversed after a number of trials, varying between 5 and 15 trials, so that the experiment consisted of a number of blocks with varying trial length (Figure 4B). Within each block, the probability of a win was fixed. The task was performed in the scanner, but here we only focus on behavioral data. 120 trials were presented for each cue (480 trials in total). Participants learned the task effectively: performance, quantified as the number of correct decisions given the true underlying probability, was significantly higher than chance across the group (t(43)=14.68, p<0.001; mean: 0.68, standard error: 0.01). The focus of the original studies using this dataset was on socio-emotional modulation of learning and choice [34,36]. Here, however, we use this dataset because the task is a probabilistic learning tasks in which the contingencies change throughout the experiment. We considered a model space including the binary VKF, binary HGF, a Kalman filter (KL) that quantifies uncertainty but not volatility, and also a Rescorla-Wagner (RW) model that does not take into account uncertainty or vary its learning rate. For all these learning models, we used a softmax rule as the response model to generate probability of choice data according to action values derived for each model as well as value-independent biases in making a go response for all these models. Note that the response model also contained value-independent biases in making or avoiding a go response due to the emotional or reinforcing content of the cues (see Methods).

**Fig 8.**
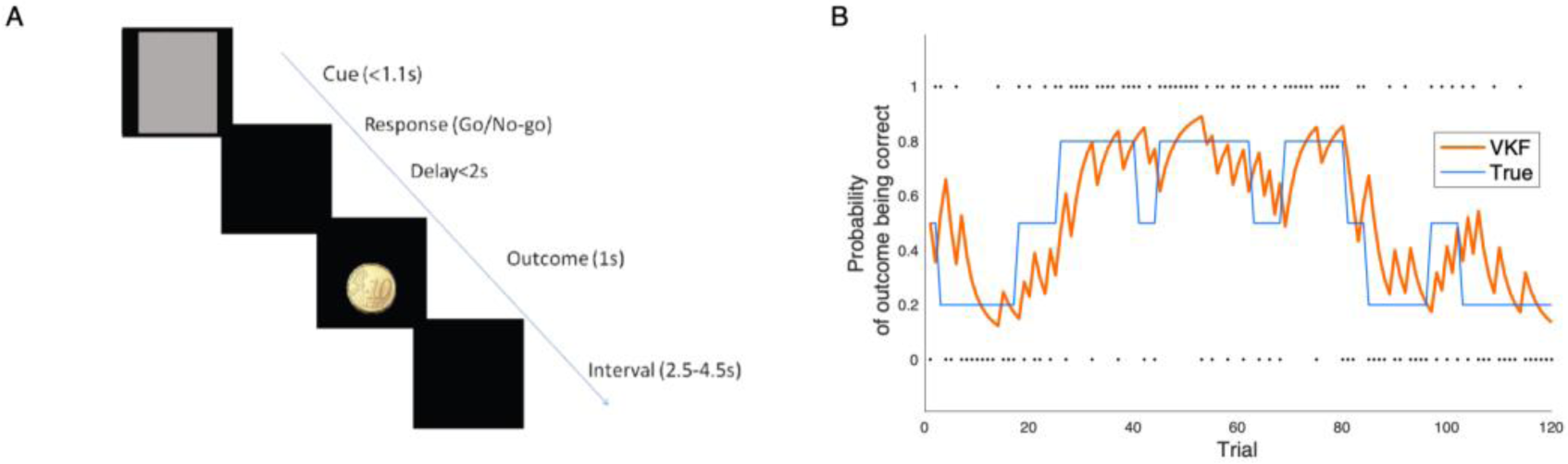
The probabilistic learning task in Experiment 1 used for testing the VKF. A) Participants had to respond (either go or no-go) after a face cue was presented. A probabilistic outcome was presented following a delay. B) An example of the probability sequence of outcome (of a go response) for one of the four trial-types and predictions of the (binary) VKF model. The response-outcome contingencies for the cues were probabilistic and manipulated independently. The dots show actual outcomes seen by the model.

This model space was then fit to choice data using HBI [35]. Importantly, the HBI combines advantages of hierarchical model for parameter estimation [37] with those of approaches that treat the model identity as a random effect [38], because it assumes that different subjects might express different models and estimates both the mixture of models and their parameters, in a single analysis. Thus, the HBI performs random effects model comparison by quantifying model evidence across the group (goodness of fit penalized by the complexity of the model [39]). This analysis revealed that, across participants, VKF was the superior model in 37 out of 44 participants (Fig 9). Furthermore, the protected exceedance probability in favor of VKF (i.e. the probability that a model is more commonly expressed than any other model in the model space [38,40] taking into account the null possibility that differences in model evidence might be due to chance [40]) was indistinguishable from 1. In a supplementary analysis, we also considered the particle filter model. Since that model is a Monte Carlo sampling method and too computationally intensive to embed within a further hierarchical estimation over subjects and other models, we fitted the model separately to each subject and compared it with the other models (see S1 Text for details). This analysis revealed that the VKF is the most parsimonious even compared to the particle filter model.

**Fig 9.**
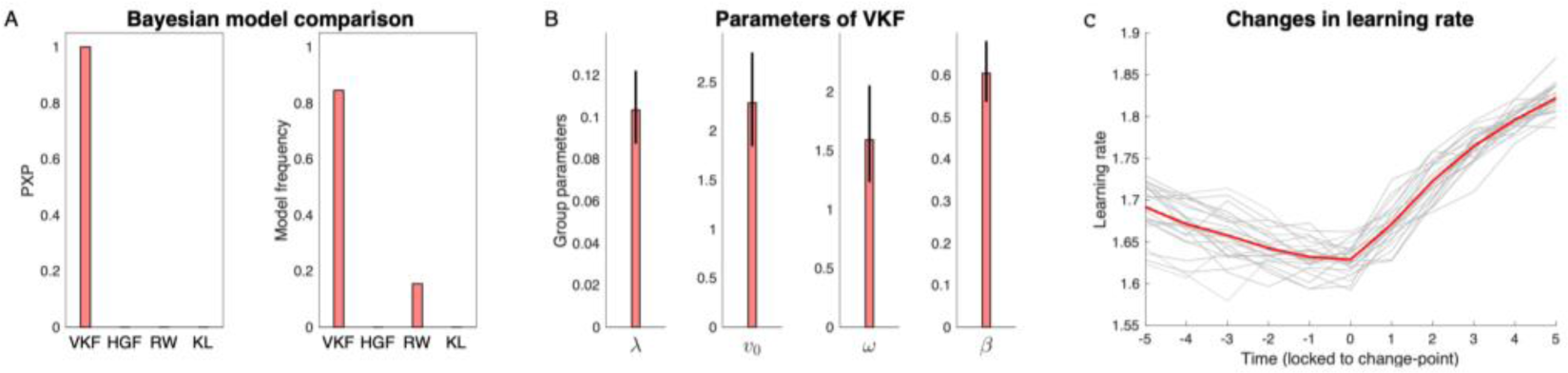
Bayesian analysis of VKF in the first experiment. A) Bayesian model comparison results comparing VKF with HGF, Rescorla-Wagner (RW) and Kalman filter (KF). Protected exceedance probabilities (PXPs) and model frequencies are reported. The PXP of the VKF is indistinguishable from 1 (and from 0 for other models) indicating that VKF is the most likely model at the group level. The model frequency metric indicates the ratio of subjects explained by each model. B) Estimated parameters of the VKF at the group-level. C) Learning rate signals, time-locked to the change points, estimated by the VKF for all participants (gray) and the mean across participants (red). The x-axis indicates trials relative to change points. The error-bars in B are obtained by applying the corresponding transformation function on the group-level error-bars obtained by the HBI [35] and, therefore, are not necessarily symmetric.

To examine the detailed behavior of the model, we also analyzed learning rate signals estimated by the VKF at the time of changes in action-outcome contingencies. For this analysis, we fitted each subject’s choice data individually to the binary VKF model to generate learning rate signals independently. Fig 9C shows variations in the learning rate signal time-locked to the change points. Across 44 participants, 40 (i.e. 91%) showed a positive change in learning rate following change points. There was a significant difference between learning rate before (obtained by averaging over 5 trials prior to change) and after change points (obtained by averaging over 5 trials) (mean increase = 0.10, P<0.001), similar to previous studies with change-point dynamics [41].

In the second experiment, 174 participants performed a learning task (originally published in [42]), in which they chose to accept or reject an opportunity to gamble on the basis of their estimation of potential reward. Thus, subjects were asked to estimate the probability of reward based on binary feedback given on every trial. The reward probability was contingent on the category of the image presented during each trial and its time-course has been manipulated throughout the experiment. During each trial, participants were also presented with the value of a successful gamble. In the original study [42], Jang et al. used this learning task, in combination with a follow-up recognition memory test, to study the influences of computational factors governing reinforcement learning on episodic memory. Here, however, we only analyze the learning part of this dataset because the task is a probabilistic learning task with switching contingencies. For every image category, the reward probability has switched at least twice during the task. Most participants learned the task effectively, as their gambling decisions varied in accordance with the probability of reward. In particular, they accepted to gamble more often in trials with high reward probability than those with low reward probability (t(173)=18.5, p<0.001; mean: 0.25, standard error: 0.01). Thirteen subjects were excluded from the analysis because a logistic regression (with intercept and two regressors: reward probability and trial value) showed a negative correlation between reward probability and their decisions to gamble, suggesting that they had not understood the task instructions. We analyzed data of the remaining 161 subjects.

We fitted the same model space used above, in which the learning models were combined with the softmax as the response model to generate probability of choices. This model space was then fit to choice data using the HBI (Fig 10). This analysis revealed that the binary VKF outperformed other models in 102 out of 161 participants (with 0.62 model frequency). Across all participants, the protected exceedance probability in favor of the VKF was indistinguishable from 1. Note that the second-best model was the Kalman filter model with model frequency of 0.3, suggesting that about 30% of subjects reduced their learning rate over time but were not sensitive to volatility. This may be because there are overall fewer switches in this task compared to the previous one. Nevertheless, across all subjects, analysis of learning rate signals time-locked to the change point shows a significant increase in learning rate (median increase = 0.03, Wilcoxon sign rank test because samples were not normally distributed, p<0.001). This effect was positive for 64% of participants.

**Fig 10.**
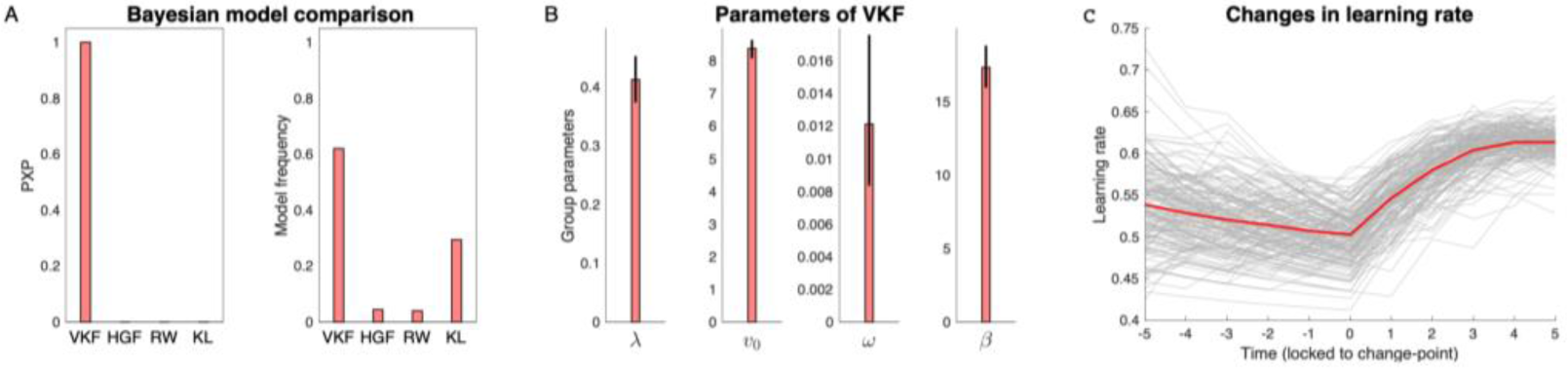
Bayesian analysis of VKF in the second experiment. A) Bayesian model comparison results comparing VKF with HGF, Rescorla-Wagner (RW) and Kalman filter (KF). Similar to the previous dataset, the PXP of the VKF is indistinguishable from 1 (and from 0 for other models) indicating that VKF is the most likely model at the group level. B) Estimated parameters of the VKF at the group-level. C) Learning rate signals, time-locked to the change points, estimated by the VKF for all participants (gray) and the mean across participants (red). The x-axis indicates trials relative to change points. The error-bars in B are obtained by applying the corresponding transformation function on the group-level error-bars obtained by the HBI [35] and, therefore, are not necessarily symmetric.

## Discussion

In this work, we have introduced a novel model for learning in volatile environments. The proposed model has theoretical advantages over existing models of learning in volatile environments, because it is based on a novel generative model of volatility that makes it possible to have a simple approximate inference model, which is also very loyal to the exact inference. Using empirical choice data in a probabilistic learning task, we showed that this model captures human behavior better than the state-of-the-art HGF.

The Kalman filter is the cornerstone of tracking theories, with widespread applications in many technological and scientific domains including psychology and neuroscience [4,7,22,23]. For example, in movement neuroscience, the Kalman filter has been used as a model of how the brain tracks sensory consequences caused by a motor command. In learning theories, the Kalman filter provides a normative foundation for selective attention among multiple conditioned stimuli predicting a target stimulus, such as food. Nevertheless, the Kalman filter is limited to environments in which the structure of process noise is constant and known. Like other models such as the HGF, the VKF fills this gap by extending the Kalman filter to inferring the process variance in volatile environments in which the variance is itself dynamically changing. In particular, VKF contains two free parameters, a volatility update rate (i.e. *λ*) indicating the extent of noise in process variance dynamics and the initial value of volatility (*v*_0_). The Kalman filter is a special case of VKF, in which the volatility update rate is zero and the constant process noise is equal to *v*_0_.

A complete understanding of a learning system requires understanding of how computational theories should be realized at the algorithmic level [17]. Previous works have shown that at this level, the normative perspective might shed light on or even encompass related psychological models, as for example temporal difference models of reinforcement learning encompass the classical Rescorla-Wagner theory. The proposed model builds a normative foundation for the intuitive hybrid models combining critical features of Rescorla-Wagner and Pearce-Hall theories for conditioning [20,21,34]. Specifically, the learning process of VKF depends on two components. The first component is the classical prediction error signal, which is defined as the difference between the observed and expected outcome, similar to Rescorla-Wagner error signal. The second component is the learning rate modulated by the volatility estimate, which is itself a function of surprise (i.e. the squared prediction error) according to Equation 13. Therefore, although the detailed algebraic computation of the surprise term slightly differs, the structure of the model is consistent with the classical Pearce-Hall model, and like it the rate of learning in VKF depends on surprise. This construction clarifies the relationship between the Pearce-Hall associability, its update, and volatility inference in hierarchical learning models such as the HGF and that of Behrens et al. [6].

The generative model of VKF is based on a novel state-space model of variance dynamics for Gaussian-distributed observations with known mean, in which the inference (for the volatility level considered in isolation) is tractable. This particular generative model leads to exact inference without resorting to any approximation. To build a fully generative model for volatile environments, we then combined this state-space model with the state-space model of the Kalman filter. Therefore, the full generative model of VKF contains two temporal chains (Fig 1), one for generating the mean and the other one for generating the variance. Although the inference is tractable within each chain, the combination of both chains makes the exact inference intractable. Therefore, we used structured variational techniques for making approximate inference, which isolates the tractable submodel as part of the variational maximization.

The state-of-the-art algorithm for learning in volatile environments is HGF [8,18], which is a flexible filter which can be extended iteratively to hierarchies of arbitrary depth. The VKF has the same conditional dependencies as the HGF with two latent levels. There are, however, two critical differences between the VKF and the HGF. First, the generative process underlying the variance is different between the two models. The HGF assumes additive Gaussian diffusion at each level, transformed by an exponential to allow it to serve as a variance for the next level down. In contrast, the generative model of process variance in the VKF uses a form of multiplicative diffusion, which guarantees non-negativity. Secondly, these generative differences lead to algorithmic ones. Although both models conduct inference under a variational approximation, due to the nonlinearity in the HGF, it is not possible to analytically maximize the variational family, and additional approximations are required. As mentioned, the generative process of the VKF is tailored to permit exact maximization of the variational distribution. Thus altogether, the VKF trades away the more general generative structure that underlies the HGF, to achieve a simpler and more accurate approximation to a more specific (two latent chains) case. We compared the performance of VKF and HGF in predicting human choice data in a probabilistic learning task, similar to those tasks that have been modeled with HGF in the recent past [8]. Bayesian model comparison showed that the VKF predicts choice data better in the majority of participants.

There is an additional difference between these models for binary outcomes, which results in a qualitative difference in the relationship of the volatility signal and learning rate (Fig 6). Consistent with classical Pearce-Hall models, surprising events increase learning rate in our model, a quality that is expected based on normative considerations [6] and empirical observations [6,32–34]. As we have shown using simulation analyses, volatility and learning rate signals are negatively correlated for binary HGF, which is a consequence of using a Taylor approximation to account for binary data. We employed a different approximation strategy to account for binary outcomes, called moment matching, which preserves the positive relationship between volatility and learning rate.

Recent work highlight the importance of uncertainty processing and its effects on learning rate for understanding a number of psychiatric disorders [33,34,43–45]. For example, Browning et al. [33] found that anxiety reduces people’s ability to adjust their learning rate according to volatility. In a recent study [34], we also found that in threatening contexts evoked by angry face images, socially anxious individuals show disruptions in benefiting from stability in the action-outcome contingency. Their dorsal anterior cingulate cortex, a region previously shown to reflect volatility estimates [6,32], also failed to reflect volatility in those contexts. This is because anxious individuals updated their expectations too rapidly in the stable conditions, possibly because they perceive any uncertainty as a signal of contingency change (i.e. volatility). Process-level models, such as VKF, can play important role in this line of research and we hope that this work be useful for quantifying critical computations underlying learning in the future.

In the last decade, scholars in the field of computational psychiatry have started to map deficits in decision making to parameters of computational models. In fact, these parameters could serve as computationally-interpretable summary statistics of decision making and learning processes. Parameters of the VKF are also particularly useful for such a mapping. There are three parameters that influence different aspects of learning dynamics in the VKF. The volatility update parameter captures the degree that the individual updates its estimate of volatility. Given the generative model of the VKF, this parameter also determines the subjective feeling of noise at the volatility level. Another parameter of the VKF is the initial volatility, which influences the learning process on early trials. These two parameters have similar effects for linear and binary observations. For the binary VKF, there is another parameter that is only relevant to the inference model (not the generative process), *ω*, which we called it the noise parameter. As shown in Fig 8, this parameter governs the scale of volatility and learning rate throughout the learning process. Notably, our simulation analyses showed that these parameters are fairly identifiable from choice data.

In this study, we assumed that the observation noise in the generative process of the VKF is a free parameter, *σ*; the HGF and traditional Kalman filters also have analogous parameters. In many situations, however, humans and other animals might have to learn the value of this noise. In particular, in addition to volatility, trial-by-trial estimation of this noise is relevant for optimal learning in situations in which observation noise might dynamically change. Deriving an efficient inference model for simultaneous tracking of both signals is substantially more difficult due to dependencies arising between variables. Our generative model of the variance and the corresponding tractable inference can be helpful for that purpose, which should be further explored in future studies.

The goal of the current study was to further process-level models of volatility by proposing a model that closely match optimal statistical inference, building on a number of studies that have been proposed in the past 15 years. Since exact inference is not possible, these studies have relied on different approximate inference approaches, such as sampling, Taylor approximation, variational inference or message-passing algorithms. Our approach for treatment of binary observations using moment matching is similar to the message-passing approach taken recently for studying dynamical systems [46,47]. We chose this approach rather than the Taylor approximation used previously [18] because it has been shown that methods based on moment matching perform better than derivative-based methods in approximating exact inference for binary Gaussian process models [48].

An important concern about Bayesian process-level models is whether their computations are biologically plausible. Similar to any other model that extends the Kalman filter, normalization is required for computing the Kalman gain in the VKF. Furthermore, our model requires the squared prediction error for updating volatility. Although performing these computations might not be straightforward with current neural network models, they are not biologically implausible. Another crucial question is how these approximate Bayesian models could be realized at the mechanistic level [49]. Recently, metaplasticity has been proposed as a mechanistic principle for learning under uncertainty [9,12,50]. Metaplasticity allows synaptic states to change without substantial changes in synaptic efficacy [51] and therefore provides a mechanism for reinforcement learning under volatility [9].

Bayesian models have recently been used for online inference of sudden changes in the environment [41,52]. Although those situations can be modeled with generative processes with discrete change-point detection, Behrens et al. [6] showed that models with volatility estimate might be as good as or even better than models with specific discrete change-points. Our simulations also showed that VKF can be successfully applied to those situations. In such situations, the volatility signal plays the role of a continuous change-point estimator, which substantially increases after a major change. This is because those sudden changes in the environments cause a large “unexpected uncertainty” signal [7,11], which substantially increases the volatility.

In this article, we introduced a novel model for learning under uncertainty. The VKF is more accurate than existing models and explains human choice data better than alternatives. This work provides new opportunities to characterize neural processes underlying decision making in uncertain environments in healthy and psychiatric conditions.

## Methods

### Simulation analysis for comparing VKF with benchmark

We implemented particle filter models using MATLAB Control Systems Toolbox. The particle filter is a sequential Monte Carlo method, which draws samples (i.e. particles) from the generative process and sequentially updates them. We implemented this model separately for the linear and binary observations, with 10000 particles and generative parameters. Note that for linear outcomes, inference on the lower level chain is analytically tractable given samples from the upper level chain. Therefore, we used RBPF [31] for the linear problem, which combines Monte Carlo sampling with analytical marginalization. We used Spearman rank correlation because signals were not normally distributed.

### Simulation analysis for comparing accuracy of VKF and HGF

The generative model of the HGF with 2 levels is based on a probabilistic model with the same dependencies as those in our generative model (Fig 1). Under this generative model, 3 chains of random variables are hierarchically organized to generate observation, *o*_*t*_:

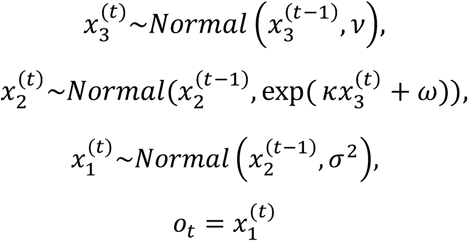

where *v* > 0 is the variance at the third level, *κ* > 0 determines the extent to which the third level affects the second level, *ω* indicates the tonic level variance at the second level, and *σ* is the observation noise. For the simulation analysis of the accuracy VKF and HGF, we generated data according to the HGF generative model (with normal observations) according to these parameters: *v* = 0.5, *κ* = 1, *ω* = −3, and *σ* = 1. The initial mean at the second and third levels were 0 and 1, respectively. We generated 1000 time-series (100 trials) using these parameters. These parameters were also then used for inference based on the HGF algorithm. However, in 82 simulations, the HGF encountered numerical problems (negative posterior variance), because its inferred trajectory conflicted with the assumptions of the inference model. For the remaining time-series, we also performed inference using an RBPF under the generative model of the HGF and the true parameters (10000 particles). The RBPF exploits the fact that inference on the lower level chain (i.e. *x*_2_) is tractable given samples from the upper level. Specifically, this approach samples *x*_3_ and marginalizes out *x*_2_ using the Kalman filter. The relative error of the HGF with respect to RBPF was defined as *E*_*HGF*_ /*E*_*RBPF*−*HGF*_ − 1, in which *E*_*HGF*_ and *E*_*RBPF*−*HGF*_ are the median of absolute differences between the ground truth generated latent variable, *x*_2_, and the predictions of the HGF and the particle filter, respectively. A similar analysis was performed by generating 1000 time-series (100 trials) using the generative model of the VKF with parameters *λ* = 0.15, *v*_0_ = 1, and *σ* = 1. We also performed inference using the RBPF based on VKF generative assumptions and the true parameters and computed the relative error for VKF, *E*_*VKF*_/*E*_*RBPF*−*VKF*_ − 1. For both algorithms, we computed correlation coefficient between their estimated signals at both levels and those from the corresponding RBPF. To compute correlation coefficient across simulations, correlation coefficients were Fisher-transformed, averaged, and transformed back to correlation space by inverse Fisher transform [53,54].

For comparing accuracy of the VKF and HGF for binary observations, we generated 500 time-series in the probabilistic switching task with binary outcomes (Fig 2). Parameters of the HGF and VKF were fitted to these time-series using a maximum-a-posteriori procedure, in which the variance over all parameters was assumed to be 15.23. The prior mean for all parameters (except *ω* in HGF) was assumed to be 0. We followed the suggestions made by Mathys et al. [55] and assumed an upper bound of 1 for both ν and *κ*. We particularly chose –1 as the initial prior mean for *ω* to ensure that the HGF is well defined at the prior mean.

### Recovery analysis of parameters

For this analysis, data were generated based on the binary VKF (Equations 14-19). In particular, the observation on trial *t, o*_*t*_, was randomly drawn based on the sigmoid-transformation of *m*_*t*−1_. The choice data were also generated randomly by applying the softmax as the response model with parameter *β*. Similar to experiment 1, for each artificial subject, we assumed 4 sequences of observations and actions (i.e. 4 cues) with 120 trials. These values were used as the group parameters: *λ* = 0.2, *v*_0_ = 5, *ω* = 1, and *β* = 1. For generating synthetic datasets for simulations, the parameters of the group of subjects (50 subjects) assigned to each model were drawn from a normal distribution with the standard deviation of 0.5.

### Implementation of models for analysis of choice data

We considered a 3-level (binary) HGF [18] for analysis of choice data, with parameters, 0 < *v* < 1, 0 < *κ* < 1, and *ω*. We also considered a constant parameter for *ω* at –4, as Iglesias et al. [8]. However, since the original HGF with a free *ω* outperformed this model (using maximum-a-posteriori estimation and random effects model comparison [38,40]), we included the original HGF in the model comparison with binary VKF. Similar to the HGF toolbox, we assumed that the initial mean at the second and third levels are 0 and 1, respectively, and the initial variance of the second and third levels are 0.1 and 1, respectively. For implementing the binary VKF, we assumed an upper bound of 10 for the initial volatility parameter, *v*_0_.

### Ethics statement

All human subjects data used here are reanalyses of anonymized data from previously published studies. Data from human subjects in experiment 1 are from a study [36] that was approved by the local ethical committee (“Comissie Mensgebonden Onderzoek” Arnhem-Nijmegen, Netherlands). Data from human subjects in experiment 2 are reported by Jang et al. [42] and the study was approved by the Brown University Institutional Review Board.

#### Experiment 1

Each of the learning models was combined with a choice model to generate probabilistic predictions of choice data. Expected values were used to calculate the probability of actions, *a*_1_ (go response) and *a*_2_ (no-go response), according to a sigmoid (softmax) function:

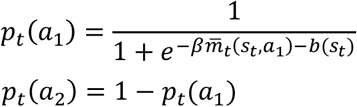

where 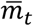was equal to *m*_*t*_ for the VKF, and it was equal to 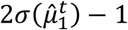 − 1 for the HGF (as implemented in the HGF toolbox). Moreover, *β* is the decision noise parameter encoding the extent to which learned contingencies affect choice (constrained to be positive) and *b*(*s*_*t*_) is the bias towards *a*_1_ due to the stimulus presented independent from learned values. The bias is defined based on three free parameters, representing bias due to the emotional content (happy or angry), *b*_*e*_, bias due to the anticipated outcome valence (reward or punishment) cued by the stimulus, *b*_*v*_, and bias due to the interaction of emotional content and outcome, *b*_*i*_. No constraint was assumed for the three bias parameters. For example, a positive value of *b*_*e*_ represents tendencies towards a go response for happy stimuli and for avoiding a go response for angry stimuli (regardless of the expected values). Similarly, a positive value of *b*_*v*_ represents a tendency towards a go-response for rewarding stimuli regardless of the expected value of the go response. Critically, we also considered the possibility of an interaction effect in bias encoded by *b*_*i*_. Therefore, the bias, *b*(*s*_*t*_), for the happy and rewarding stimulus is *b*_*e*_ + *b*_*v*_ + *b*_*i*_, the bias for the angry and punishing stimulus is −*b*_*e*_ − *b*_*v*_ + *b*_*i*_, the bias for the happy and punishing stimulus is *b*_*e*_ − *b*_*v*_ − *b*_*i*_ and the bias for the angry and rewarding stimulus is −*b*_*e*_ + *b*_*v*_ − *b*_*i*_.

#### Experiment 2

This experiment was conducted by Jang et al. [42] to test the effects of computational signals governing reinforcement learning on episodic memory. The learning task consisted of 160 trials. On each trial, the trial value was first presented, followed by the image (from one of the animate or inanimate categories), response (play or pass) and the feedback. The feedback was contingent on the response made by the participant and given the probability of reward given the image category. Thus, if the participant chose play and the trial was rewarding, they were rewarded the amount shown as the trial value. If they chose to play and the trial was not rewarding, they lost 10 points. If the choice was pass, the participant did not earn any reward (i.e. 0 point), but was shown the hypothetical reward of choosing play. Data were collected using Amazon Mechanical Turk.

For modeling choice data, the softmax function with parameter *β* was used as the response model, in which the expected value of play was calculated based on the probability of reward estimated by the learning models and the value of trial shown at the beginning of each trial. The expected values were divided by 100 (the maximum trial value in the task) before being fed to the softmax to avoid numerical problems.

### Model fitting and comparison

We used a hierarchical and Bayesian inference method, HBI [35], to fit models to choice data. The HBI performs concurrent model fitting and model comparison with the advantage that fits to individual subjects are constrained according to hierarchical priors based on the group-level statistics (i.e. empirical priors). Furthermore, the HBI takes a random effects approach to both parameter estimation and model comparison and calculates both the group-level statistics and model evidence of a model based on responsibility parameters (i.e. the posterior probability that the model explains each subject’s choice data). The HBI quantifies the protected exceedance probability and model frequency of each model, as well as the group-level mean parameters and corresponding hierarchical errors. This method fits parameters in the infinite real-space and transformed them to obtain *actual* parameters fed to the models. Appropriate transform functions were used for this purpose: the sigmoid function to transform parameters bounded in the unit range or with an upper bound and the exponential function to transform parameters bounded to positive value. To ensure that the HGF is well defined at the initial prior mean of fitted parameters (i.e. zero), we assumed *ω* = *ω*_*f*_ − 1, where *ω*_*f*_ is the fitted parameter.

The initial parameters of all models were obtained using maximum-a-posteriori procedure, in which the initial prior mean and variance for all parameters were assumed to be 0 and 6.25, respectively. This initial variance is chosen to ensure that the parameters could vary in a wide range with no substantial effect of prior. These parameters were then used to initialize the HBI. The HBI algorithm is available online at https://github.com/payampiray/cbm.

## Acknowledgment

We are grateful to Matthew Nassar for sharing the empirical dataset used in this study.

## Supplementary Materials

### Control analyses for comparison between VKF and HGF

We first verified that although the two generative models are parameterized differently, their parameters models were chosen in comparable regimes with respect to the ultimate inference problem of tracking the latent state *xt*. In particular, the median trial-by-trial change in that lower-level signal, which is defined similarly in both models based on a Gaussian random walk, was comparable. The average of this measure across all simulations for HGF and VKF was 1.84 and 1.93, respectively.

Performance of the models might depend on the parameters determining variability of the volatility signal. In the HGF and VKF, this depends on *ν* and *λ*, respectively. Therefore, we performed a second analysis in a different parameter regime by reducing *ν* from 0.5 (the original analysis) to 0.25 for the HGF, and by reducing *λ* from 0.15 (original analysis) to 0.1 for the VKF. The relative error of the HGF and VKF was 8.20% (SE=0.6%) and 3.2% (SE=0.3%), respectively. The measure of trial-by-trial changes in the lower-level signal, as defined above, for the HGF and VKF was 0.68 and 1.01, respectively, indicating that the parameters generate comparable signals, and if anything, the tracking problem for VKF is a bit harder in that the latent variable is a bit less stable.

Next, we considered the possibility that differences between the models’ relative performance scores might arise not due to their particular inferential approximations, but instead due to the particle filtering algorithm used as a baseline. In particular, while it is important to use such a baseline, so as to measure performance relative to a measure of what is optimally achievable in a particular generative family, actual optimal inference is intractable, so we rely on the RBPF as a proxy. However, that algorithm is also approximate (though computationally intensive and with good convergence properties), and in principle its error might also differ between the generative processes. Accordingly, we also compared the two models relative to a second baseline independent of (and algorithmically simpler than) the RBPF, this one an idealistic baseline provided by an ideal Kalman filter augmented with an (unrealistic) oracle providing the true volatility state at each step. This construction exploits the fact that both VKF and HGF reduce to the Kalman filter given the volatility. In particular, for every timeseries generated in the original analysis, we simulated the Kalman filter in which the volatility parameter on every trial was given by the true volatility. Note that this is not a realistic inference model, as it has access to information (i.e. true volatility) that other inference models do not. This idealistic Kalman filter model was then used as the baseline to obtain a measure of relative error for both the VKF and HGF, similar to the previous analysis. Across all simulations, the relative error for the VKF and HGF was 5.5% (SE=0.4%) and 46.6% (SE=6.7%), respectively, indicating that the inference by the VKF was closer to the idealistic Kalman filter.

All these results comparing accuracy of VKF and HGF in inference on timeseries generated by their own generative models for both sets of parameters per model are summarized in the following table.

|  |  | Number of conflicts | Relative error– RBPF | Relative error– ideal Kalman | Correlation coefficient state | Correlation coefficient volatility |
| --- | --- | --- | --- | --- | --- | --- |
| Set 1 | HGF | 82 | 21.4% | 46.6% | 1.00 | 0.88 |
|  | VKF | NA | 2.7% | 5.5% | 1.00 | 0.95 |
| Set 2 | HGF | 15 | 8.2% | 25.8% | 1.00 | 0.88 |
|  | VKF | NA | 3.2% | 5.0% | 1.00 | 0.85 |

### Control analyses for testing effects of volatility on learning rate for binary outcomes

We performed a control analysis to ensure that positive and negative relationship between volatility and learning rate is a general behavior of binary VKF and binary HGF, respectively. We simulated these models with randomly drawn parameters (1000 simulations) using a single sequence of binary outcomes (those plotted in Fig 6). The volatility and learning rate signals were positively correlated for binary VKF in all simulations, whereas those were negatively correlated in 94% of simulations for binary HGF.

### Subject-level fitting of models to empirical data

In addition to the HBI hierarchical fitting approach discussed in the main text, to verify the generality of the result, we also fit empirical data to the models at the individual per-subject level using a Laplace approximation. For the four models reported in the main text, this is equivalent to the first iteration of the HBI. We also included the particle filter (PF) model in this analysis using a simple sampling procedure (1000 samples) to first draw generative parameters of the VKF (i.e. *v*0 and *λ*), fit the PF (10000 particles) given those parameters to obtain trial-by-trial predictions, and then fitted the parameters of the response model using the Laplace approximation. To quantify model evidence at the subject-level, we used Bayesian information criteria (BIC) to account for the generative parameters. The following table shows the results for both experiments in terms of model frequency (MF) and protected exceedance probability (PXP).

| Model | Experiment 1 |  | Experiment 2 |  |
| --- | --- | --- | --- | --- |
|  | PXP | MF | PXP | MF |
| VKF | 0.85 | 0.40 | 1 | 0.76 |
| HGF | 0.01 | 0.12 | 0 | 0.05 |
| RW | 0.09 | 0.24 | 0 | 0.04 |
| KF | 0.04 | 0.21 | 0 | 0.15 |
| PF | 0.01 | 0.02 | 0 | 0.01 |

## Supplementary Appendix

### Appendix A

In this appendix, we give a formal treatment of the VKF. We first review state space models and principles of making inference in those models. Next, we explain two specific state space models with different types of noise. The first one is Kalman filter, a well-known model in which the noise is additive and Gaussian. Next, we present another space model, based on a class of non-Gaussian state space models, in which the noise is multiplicative. Inference for these models is tractable given some assumptions about the noise. Building on these two tractable models, we are then in the position to provide an approximate solution for inference in volatile environments by using the powerful technique of structured variational inference. This technique makes use of exact inference algorithms on tractable substructures of an intractable model and provides an approximate variational solution.

#### A.1 State Space Models

We begin by describing state space model, in which a sequence of observations, {*u*_*i*_} where *i* = 1, …, *t*, is modeled by specifying a probabilistic relation between the observations and a sequence of latent states, {*s*_*i*_} and a Markov structure linking the latent states. A state space model assumes that i) *u*_*t*_ is independent of all other observations and states given *s*_*t*_, and ii) *s*_*t*_ is independent of all other states given *s*_*t*−1_. Thus, the joint distribution for the state space model is given by

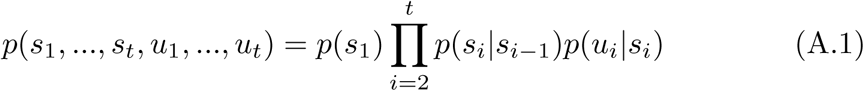

The inference goal in the state space model is to obtain the posterior over *s*_*t*_ given observations. It can be shown then that regardless of the form of transition probabilities and the probabilistic relation between *s*_*t*_ and *u*_*t*_, the posterior can be obtained recursively. First, the current state estimate is projected ahead in time. In other words, the posterior of *s*_*t*_ given all observations prior to *t* is calculated based on the previous posterior *q*(*s*_*t*−1_) = *p*(*s*_*t*−1_|*u*_1_, …*u*_*t*−1_):

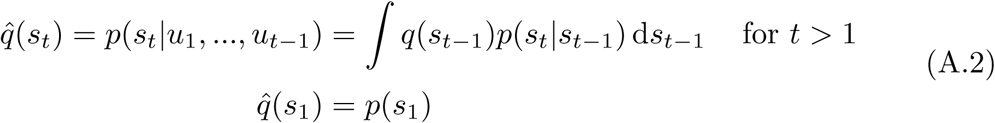

Next, the projected estimate is adjusted by actual measurement, giving rise to the posterior given all observations:

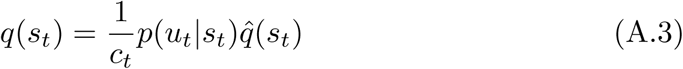

where *c*_*t*_ = *p*(*u*_*t*_|*u*_1_, …, *u*_*t*−1_) is constant with respect to *s*_*t*_. Note that since *c*_*t*_ is independent of *s*_*t*_, it is only needed to be calculated if one is interested in the likelihood of data, which is easy to see to be given by

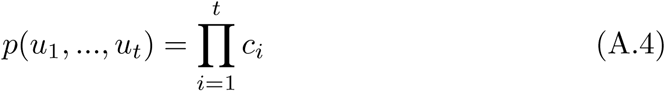

Furthermore, the posterior over two consecutive states *s*_*t*−1_ and *s*_*t*_ is given by

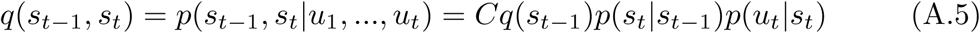

where *C* is a normalization constant independent of *s*_*t*−1_ and *s*_*t*_.

As noted above, equations (A.2-A.5) are valid for any model with conditional dependencies as given in equation (A.1) regardless of the form of *p*(*s*_*t*_|*s*_*t*−1_) and *p*(*u*_*t*_|*s*_*t*_). However, the form of *p*(*s*_*t*_|*s*_*t*−1_) and *p*(*u*_*t*_|*s*_*t*_) determine whether the integral in equation (A.2) is tractable and whether the process can be summarized in an efficient and simple algorithm by ensuring that *q*(*s*_*t*_) has the same functional form as *q*(*s*_*t*−1_). Kalman filter provides a tractable solution for a state space model in which transition probabilities are Gaussian with a fixed variance, which is suitable for prediction in stable environments. We discuss Kalman filter in section A.2. In section A.3, we build another state space model in which observations are Gaussian with a stochastic variance determined by the states. Furthermore, while the source of stochasticity in state transitions of Kalman filter is additive noise, the source of stochasticity in the second state space model is a multiplicative noise, which makes it appealing as a model of uncertainty. Then, in section A.4, we combine the two models to address prediction problems in volatile environments.

#### A.2 Kalman filter

Consider a state space model in which, on trial *n*, hidden state of the environment, *x*_*t*_, is equal to its state on trial *n* − 1 plus some Gaussian noise.

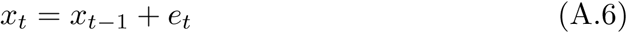

where *e*_*t*_ is a zero-mean Gaussian random variable with variance *v*. Therefore, the generative probabilistic model of *x*_*t*_ is

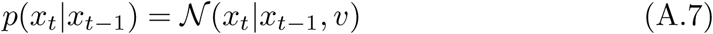

The initial state is also assumed to be Gaussian

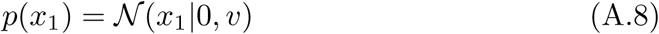

Observations are given by a Gaussian distribution:

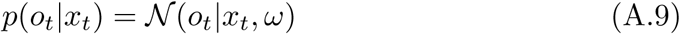

where *ω* is the observation variance. Using equations (A.2-A.3), we obtain:

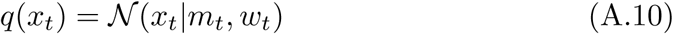

where

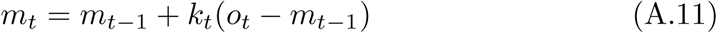

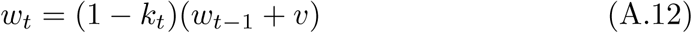

and *k*_*t*_ is the learning rate (also called Kalman gain) and is given by:

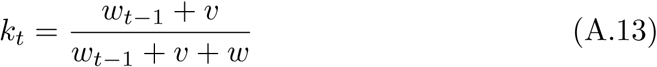

Furthermore, we can make use of (A.5) to obtain the covariance between consecutive states, *x*_*t*−1_ and *x*_*t*_:

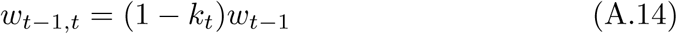

#### A.3 A tractable filter for variance

Let’s now consider another state space model in which hidden state on trial *t, z*_*t*_, depends on its value on previous trial, *z*_*t*−1_ multiplied by some (independent) noise, *ϵ*_*t*_. Then, a Gaussian random variable, *x*_*t*_, is drawn with a known mean, *µ*_*t*_, and a precision given by *z*_*t*_,

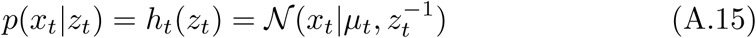

Therefore, conditional dependencies indicate that equations (A.2-A.3) hold for the posterior over *z*_*t*_.

Our goal in this section is to build a rich model of dynamic (inverse) variance with tractable and simple update equations. First, we inspect the functional form of *h*_*t*_(*z*_*t*_) with respect to *z*_*t*_:

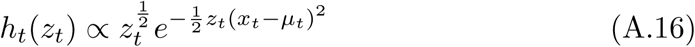

Therefore, *h*_*t*_(*z*_*t*_) takes a form of gamma distribution in which information about new observations is always coming through the rate and the shape is constant. Moreover, equation (A.3) indicates that if 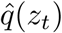 is a gamma distribution, then *q*(*z*_*t*_) takes the form of gamma distribution. Thus, for the moment, we assume that *q*(*z*_*t*_) is a gamma distribution with a constant shape *a* and a dynamic rate *b*_*t*_:

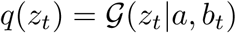

In the previous section, we saw that the transition noise was additive. For variance, it makes sense to assume a multiplicative noise. Thus, we suppose that the value of *z*_*t*_ is equal to *z*_*t*−1_ multiplied by some noise

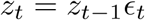

Assuming that the noise is bounded between zero and *R ≥* 1, we can write this equation as

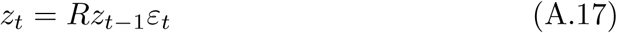

where *ϵ*_*t*_ = *Rε*_*t*_, in which *ε*_*t*_ is constrained in the unit range and has a beta distribution. Assuming that conditional expectation of *z*_*t*_ is *z*_*t*−1_, we obtain 𝔼[*ϵ*_*t*_] = *R*^−1^. A general beta distribution with this property is given by:

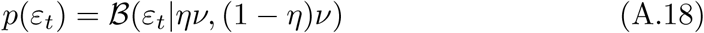

where *η* = *R*^−1^ and *ν* is a constant positive parameter. We show below that there is a specific value for *ν*, which makes the inference tractable.

Using transform theorem for random variables, the conditional distribution of *z*_*t*_ is given by

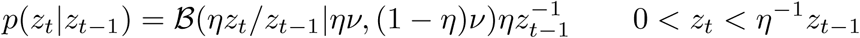

Applying equation (A.2) to 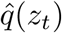, we obtain

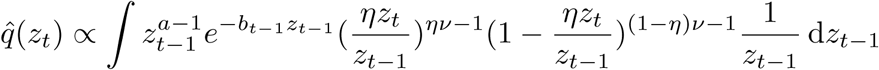

If we take *a* to be equal to *ν*, the integral is tractable resulting in a gamma distribution for 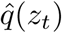. Note that evaluating those terms that are independent of *z*_*t*_ is not even necessary because 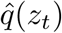 is a probability distribution. Therefore, we have

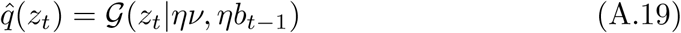

Combining 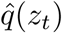 with *h*(*z*_*t*_), we obtain *q*(*z*_*t*_):

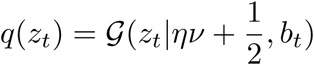

where

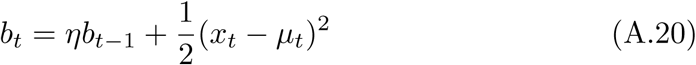

If we now take the shape of *q*(*z*_*t*_) to be the same as the shape of *q*(*z*_*t*−1_), the inference is similarly tractable for all next trials. Consequently, *ν* is given by

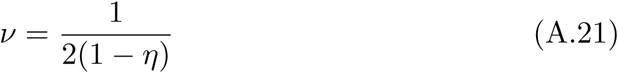

Therefore, the posterior distribution is a gamma distribution with a fixed shape and an evolving rate

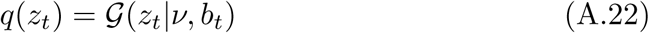

We can also write *b*_*t*_ in terms of *v*_*t*_ = 𝔼[*z*_*t*_]^−1^

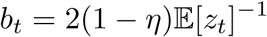

Now we can also write (A.20) in terms of *v*_*t*_ = 𝔼[*z*_*t*_]^−1^

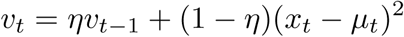

If we define *λ* = 1 − *η*, we can write this equation in the form of an error-correcting update rule:

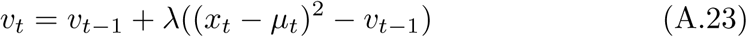

It is also important to note that 𝔼[*z*_*t*_] under 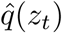 is given by *v*_*t*−1_ according to equation (A.19). Furthermore, as equation (A.2) indicates 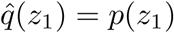, we take the probability of initial state to be given by

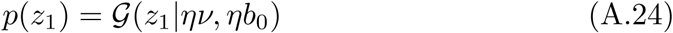

where *b*_0_ *>* 0 is a free parameter, which we can equivalently write it according to the initial volatility value *v*_0_. Note that as a consequence of making the inference tractable, the form of multiplicative transition noise is restricted according to equation (A.21). However, this is not a major restriction, as the mean of noise is still free to be chosen. Note that it is also possible to assume other forms for *ν*, although those assumptions might make the inference slightly more complicated.

#### A.4 Volatile Kalman filter

In this section, we provide a solution to the general prediction problem in volatile environments. Specifically, consider a problem in which on trial *t* a latent random variable, *x*_*t*_, is given by its previous value, *x*_*t*−1_, plus some Gaussian noise with a precision given by another dynamic random variable, *z*_*t*_:

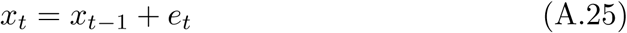

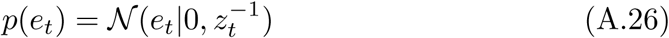

The dynamic of *z*_*t*_ is given by equations (A.17-A.18). The observation on trial *t, o*_*t*_, is then drawn based on the value of *x*_*t*_ according to a normal distribution:

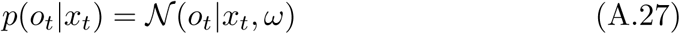

Therefore, the graphical model corresponding to this problem consists of two coupled chains of latent variables, *x*_*t*_ and *z*_*t*_. Exact inference for this problem is not possible due to coupling between the two chains. However, it is possible to derive a variational approximation by inducing factorization between the two chains, while preserving all dependencies within each chain. Here, we assume

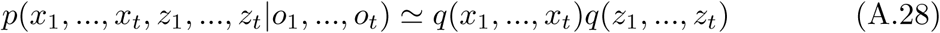

Using variational calculations, we obtain

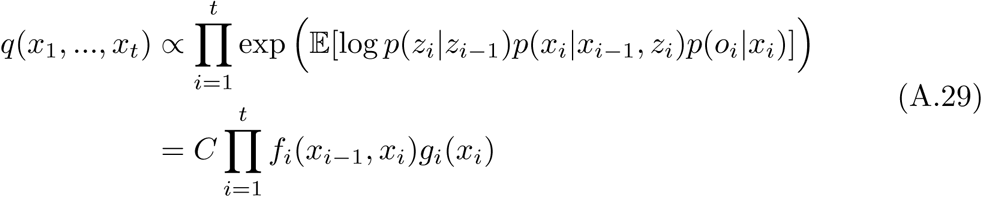

where all terms independent of *x*_1_, …, *x*_*t*_ are absorbed into *C* and

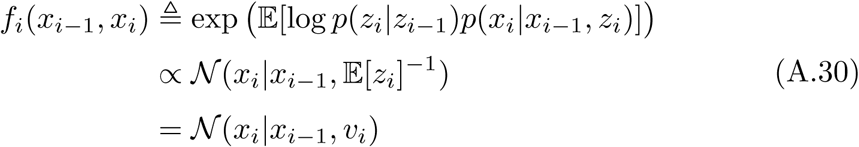

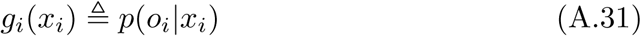

Since the conditional dependencies for *x*_*i*−1_ and *x*_*i*_ in (A.29) is the same as those in (A.1), exact inference for *q*(*x*_*t*_) follows (A.2-A.3). Moreover, since dependencies are in Gaussian forms, we obtain *q*(*x*_*t*_) = *𝒩*(*x*_*t*_|*m*_*t*_, *w*_*t*_), we can use Kalman equations (A.10-A.13) to update *m*_*t*_ and *w*_*t*_ in which we replace *v* in those equations with *v*_*t*_.

Next, we use variational techniques to obtain *q*(*z*_*t*_):

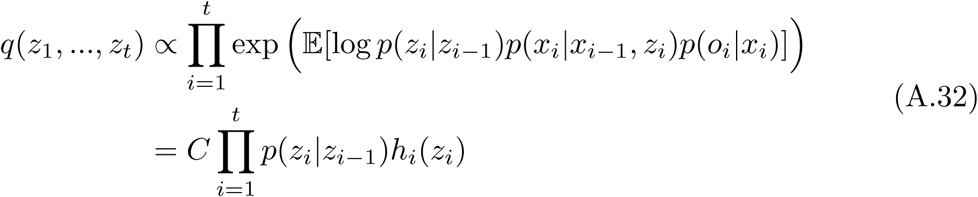

where all terms independent of *z*_1_, …, *z*_*t*_ are absorbed into *C* and

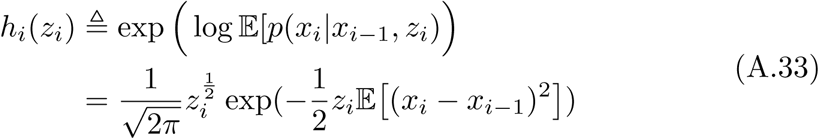

Since the conditional dependencies for *z*_*i*−1_ and *z*_*i*_ in (A.32) is the same as those in (A.1), inference for *q*(*z*_*t*_) follows (A.2-A.3). Moreover, since dependencies between *z*_*i*−1_ and *z*_*i*_ have the same functional form as (A.16) and (A.17), the posterior *q*(*z*_*t*_) takes the form of Gamma distribution, in which its mean, *v*_*t*_, gets updated according to 𝔼 [(*x*_*t*_ − *x*_*t*−1_)^2^]:

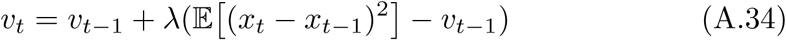

Since the expectation in (A.34) should be taken under the posterior *q*(*x*_*t*−1_, *x*_*t*_), we have

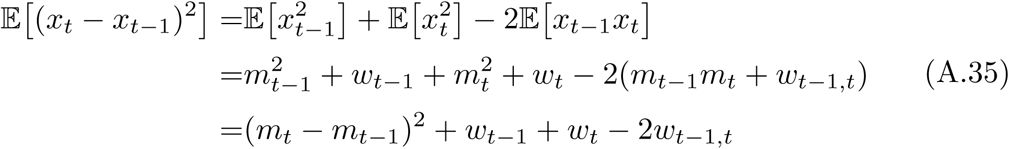

These results give rise to an efficient and simple algorithm for learning in volatile environments. On every trial *t*, we first compute 𝔼[*z*_*t*_]^−1^ under 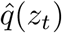, which is given by *v*_*t*−1_. Next, we update *m*_*t*_ and *w*_*t*_ and *v*_*t*_, according to equations (A.11-A.14) by replacing *v* with 𝔼[*z*_*t*_]^−1^ = *v*_*t*−1_. Finally, making use of equations (A.35), (A.12) and (A.14), we can simplify (A.34) and obtain equation (8). Note that here, unlike typical algorithms obtained based on variational Bayes, we only update statistics once to have an efficient algorithm for learning.

#### A.5 Binary VKF

In this appendix, we give a formal treatment of the binary VKF, in which observations are drawn using the sigmoid function:

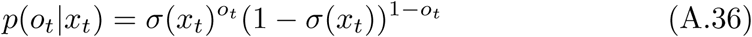

where *σ*(.) is the sigmoid function:

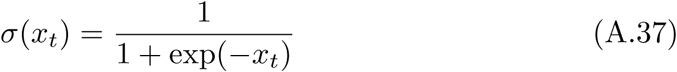

Note that for binary observations, (A.3) indicates that *q*(*x*_*t*_) is not normally distributed even if we assume that 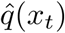 is normal. Therefore, some approximate strategies are required for binary observations.

A well-known method approximates *q*(*x*_*t*_) by matching its moments to those of 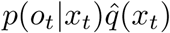. For a Gaussian distribution, this means that first and second order moments should be matched. However, since matching variance might lead to negative values for the variance, we assume that the variance is constant (similar to Kalman filter) and match only the mean of distributions. Formally, we take the “message” from *o*_*t*_ to *x*_*t*_ to be an unnormalized Gaussian, 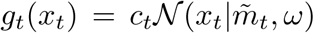, in which the variance *ω* is a constant and the mean will be obtained by matching the mean of *q*(*x*_*t*_) = *𝒩*(*x*_*t*_|*m*_*t*_, *w*_*t*_) with that of 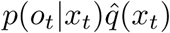, in which 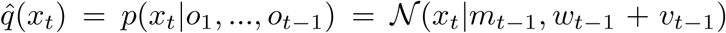. We used a simple approximation for the arising integral, which is inspired by Mackay (1992):

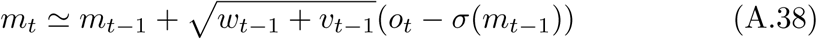

It is also possible to approximate *m*_*t*_ using a slightly more complicated equation given by MacKay (1992). Using simulation analyses, however, we found that (A.38) results in better approximations. The variance of *q*(*x*_*t*_) is then given by

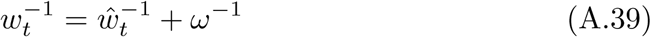

which results in equations (6-8).

